# How many crystal structures do you need to trust your docking results?

**DOI:** 10.1101/2025.09.19.677428

**Authors:** Alexander Matthew Payne, Benjamin Kaminow, Hugo I. MacDermott-Opeskin, Iván Pulido, Jenke Scheen, Maria A Castellanos, Daren Fearon, John D. Chodera, Sukrit Singh

## Abstract

Structure-based drug discovery relies on the prediction of protein-bound poses of new molecule designs, the accuracy of which can impact downstream prioritization. While it is expected that crystal structures of similar molecules would provide the best template for predicting the poses of new designs, the time and cost required motivates identifying a point of diminishing returns for collecting new structures. Using 403 crystal structures of SARS-CoV-2 main protease from the open science COVID Moonshot project, we explore the tradeoff between the cost and utility of obtaining crystal structures for accurately predicting poses of designed molecules. We observe that similar reference ligands enable superior pose prediction and show that success plateaus after approximately five crystal structures per generic Bemis-Murcko scaffold, exceeding 95% for the campaign’s lead series. This work provides practical recommendations for resource allocation in structure-enabled drug discovery campaigns.

**Table of Contents Graphic:** 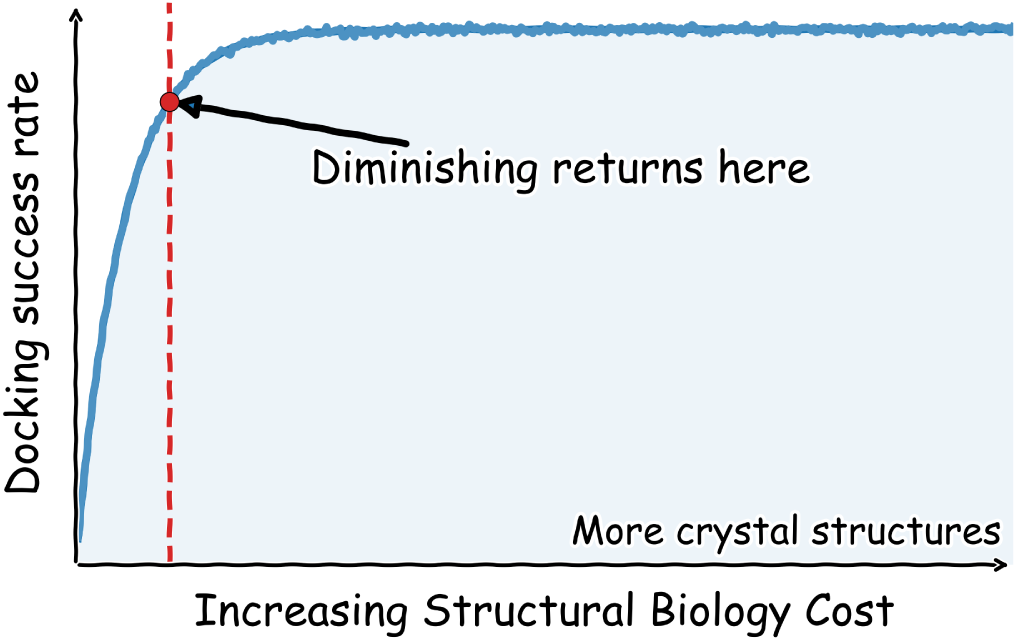

## Introduction

### Structure-based drug discovery has accelerated the development of new therapeutics

Structure-based drug discovery (SBDD) is an established paradigm for leveraging the wealth of structural information to accelerate the development of new therapeutics.^1–4^ SBDD is founded on the idea that knowledge of the structure of a drug target—and how an early hit or lead molecule binds that target—can accelerate the rational design of potent and selective compounds on the way to preclinical development (Figure 1). The effectiveness of this acceleration is difficult to assess, as it is cost-prohibitive to run the same program in parallel with and without structural elucidation experiments. SBDD continues to become more popular since early applications to the development of HIV protease inhibitors; ^2,5–7^ as of 2023, at least 65% of successful hit-to-clinical progressions used SBDD^8^ with 33% of programs from 2015–2022 using SBDD for hit-finding.^9^ As computational models improve their ability to predict binding modes and translate them into affinity or selectivity readouts, SBDD approaches can rapidly decrease time toward potency and selectivity goals in fewer iterative cycles (Figure 1).^4,5^

**Figure 1:**
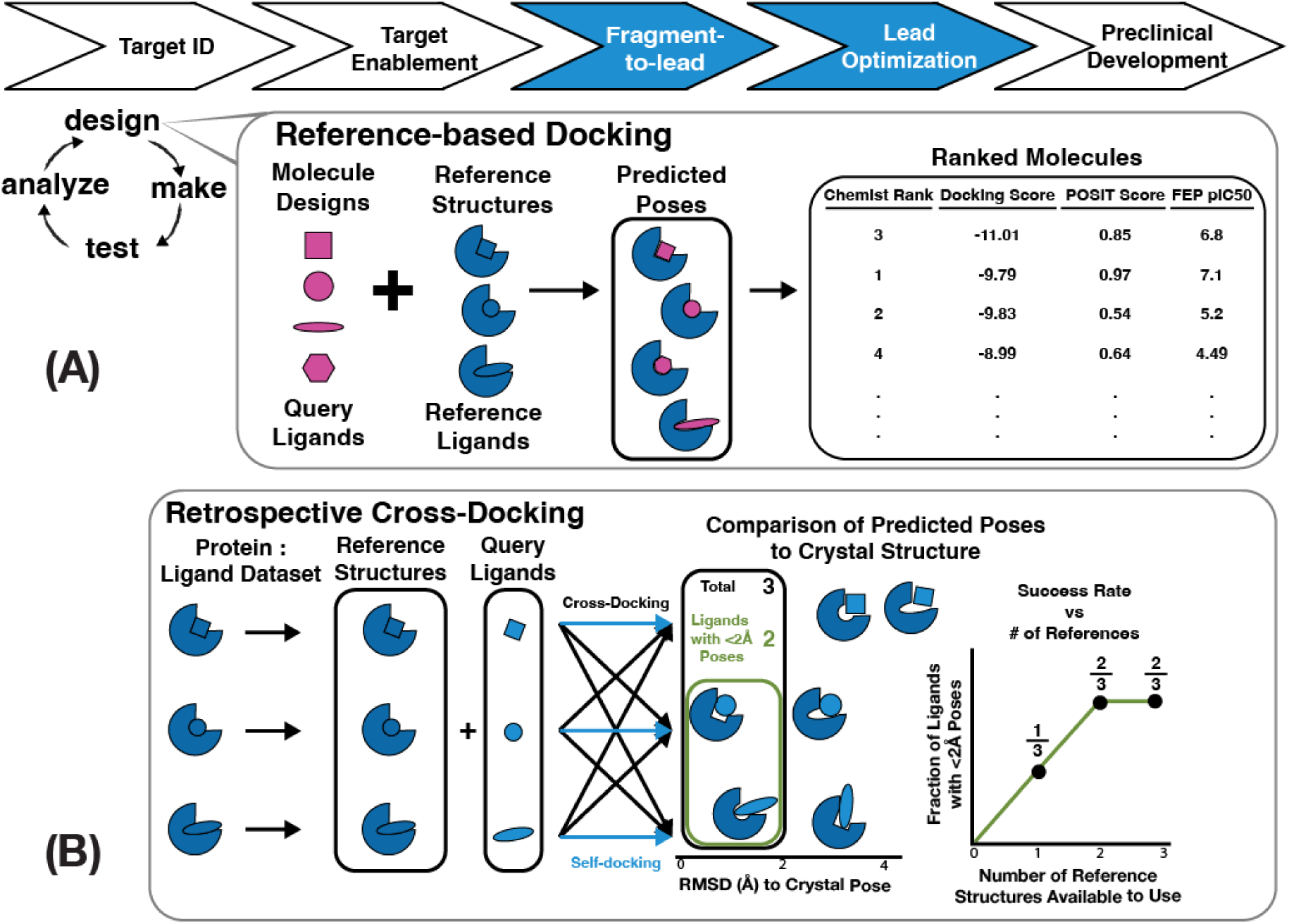
Retrospective cross-docking quantifies how pose-prediction performance depends on the number and nature of available structures. (A) Pose prediction is a critical component of the fragment-to-lead and lead-optimization stages of structure-based discovery programs (top, blue) during the design phase of the design-make-test-analyze (DMTA) cycle (left). Reference-based docking (inset) takes advantage of existing structures with reference ligands bound to the target (inset, blue) in order to pose the query ligands (inset, pink). Posed query molecules can be evaluated in multiple ways (inset, right), including structure-activity relationship rationalization by medicinal chemists, virtual screening using docking and consensus scoring functions, and binding affinity predictions using alchemical free energy calculations. (B) Retrospective cross-docking enables a thorough investigation of pose-prediction success rate. In this study, we evaluate the success rate of pose prediction as the fraction of ligands with a pose ≤ 2 Å from its experimental structure (green box). Success rate is measured for different methods for ranking the predicted poses (right).

### The development of computational methods that use protein structures has increased the predictive power of structure-based drug design

Early SBDD efforts mainly used structures to qualitatively guide design by rationalizing the observed structure-activity relationship (SAR) of a ligand series (usually congeneric).^10^ At present, many computational methods exist that use protein-ligand structures for quantitative predictions in the broader field of computer-aided drug discovery (CADD).^11^ From empirical high-throughput virtual screening^12–14^ to physics-based simulations for binding affinity prediction^15–20^ to machine-learning models for generative chemistry,^21–24^ the toolbox of computational methods has never been larger or more diverse. Over the last five years, deep-learning methods have increasingly been applied in structure-based drug design for protein structure prediction,^25–28^ quantum mechanical functionals approximation,^29–31^ and even directing the use of these methods with active learning.^32–34^

Historically, docking, which uses a mix of physics-based and empirical scoring functions to evaluate ligand poses, has driven prediction of ligand binding modes for a congeneric series.^35–37^ This approach has become a key component of modern high-throughput virtual-screening.^14,36^ More recently, ligand-protein co-folding approaches have emerged as an alternative, though there is no standard way yet to enforce that these methods model ligands in the same binding mode as a reference structure.^35–42^ Current pose-prediction benchmarks have not comprehensively assessed the ability of co-folding methods to consistently and prospectively predict binding modes. As such, experimental structural validation continues to play a critical role in SBDD.

### Improvements in high-throughput X-ray crystallography have increased the feasibility for structure-driven drug discovery

Recent experimental advances have greatly decreased the complexity of structurally enabling a drug discovery project.^6,43,44^ Development of automated, make-on-demand chemistry, fragment-based drug discovery (FBDD), and improvements in beamline engineering and analysis have made it possible to collect hundreds to thousands of fragment-bound structures in weeks.^44–46^ Organizations such as the Diamond Light Source have pioneered these efforts, resulting in many new datasets of protein-ligand structures for therapeutically relevant targets.^47–53^

Despite substantial advances, crystallography remains a costly and time-consuming endeavor with potential bottlenecks throughout the process. Producing a protein sample that is both biochemically relevant and crystallizable is often a major hurdle for each new project. Even once obtained, identifying crystallization conditions that yield suitable crystals introduces another layer of unpredictability. Added to this are the logistical challenges of securing regular beamtime and shipping samples to synchotron facilities. These barriers are well recognized in the SBDD community and raise the central questions we address this in this work:^54–56^ *How many structures are needed to drive progress with computational models? Can we define a point of diminishing returns, or is more structural data always better?*

### Retrospective cross-docking enables evaluation of reference-based pose prediction

As accurate pose prediction is foundational to SBDD efforts, we evaluate the gained utility of additional crystal structures by examining how available structures affect pose prediction.^57^ We distinguish pose-prediction as its own task from virtual screening, which typically refers to selecting a subset of molecules from a database of candidates for testing in a downstream assay, whereas pose-prediction refers to an *in-silico* determination of the 3D coordinates of a protein-ligand complex for a selected ligand. Ligand-based virtual screening assesses whether proposed candidates are likely to bind to the target using only molecular properties of the ligand. Structure-based virtual-screening takes advantage of SBDD by both predicting the pose of the ligand in the target and using a scoring function to rank the posed molecules. Pose-prediction was first developed as the DOCK software in 1982,^58^ and several of these early docking algorithms are still in use today.^59–61^ Algorithmic^62^ and scoring^63,64^ improvements have increased the speed, versatility, and accuracy of these programs. One of the early examples of ‘reference-based’ docking, shown by OpenEye’s HYBRID method, improves traditional (protein-only) docking algorithms (OpenEye FRED) by combining structure-based and ligand-based scoring functions (hybrid docking). HYBRID docking uses a modified scoring algorithm that prioritizes query ligand poses which maximize the overlap with the crystallographic ligand. ^65,66^ OpenEye’s POSIT algorithm improves upon hybrid docking by evaluating poses using the POSIT Probability, an estimate of the likelihood that the ligand will bind within 2 Å of the predicted pose.^67^ OpenEye’s pose prediction methods are readily available for academic testing and benchmarking, providing an industry-standard suite to assess pose-prediction.

Reference-based (also called templated, or template-based) docking is primarily useful during the hit-to-lead (or fragment-to-lead, in FBDD) and lead optimization phases of the drug discovery pipeline when crystal structures are available (Figure 1A). Pose prediction models are usually evaluated in self-docking experiments, where ligands from protein-ligand crystal structures are redocked into their corresponding crystal structures.^66,68^ Performance is evaluated by computing the heavy-atom root-mean-squared deviation (RMSD) of the predicted poses to their crystallographic poses.^66,68^ Traditional pose prediction has considered an RMSD *≤* 2 Å as a sufficiently accurate binding pose, in part due to the resolution ranges found in many ligand-bound crystals.^61^ Although self-docking studies benefit from datasets with only a single structure per ligand, they cannot evaluate cases in which structures of related ligands are available. Cross-docking on the other hand, mimics real-world scenarios by docking each ligand from a dataset of protein-ligand structures to every other protein structure, enabling a comprehensive analysis of pose prediction performance (Figure 1B). Few publicly available datasets that reflect a real hit-to-lead optimization pipeline (”lead-opt” datasets) are rich in both binding affinities and crystal structures. As an exception, kinase domains have been abundant in public datasets as they readily crystallize and are a thoroughly-mined target for drug discovery given their prevalence in many therapeutic areas.^69,70^ The Volkamer Lab recently ran a retrospective cross-docking analysis of over 423 kinase inhibitors bound to 10 different kinases.^71^ They found that POSIT outperforms FRED for similar reference and query molecules, and that there are diminishing returns in performance gain after 20-50 structures.^71^ Temporally split cross-docking has previously been used as an approach to evaluate prospective performance.^72^ The PINC benchmark, consisting of ten targets and 949 ligands, predicts ligand structures solved after a specific cutoff date using only structures available prior to the cutoff. The use of the PINC benchmark finds that similarity-guided reference selection raises success rates 20–30% to 60% for the top-ranked pose family. Template-based approaches have also seen further development: FitDock fits an initial conformation to a template by hierarchical feature alignment and improves success by 40–60% over template-free docking with AutoDock Vina.^73^ Such prior approaches and studies establish that template information improves cross-docking and a temporal split is an appropriate way to assess performance. Each previous work focused on a few structures per target and thus it remains unanswered whether pose-prediction is dependent on the number of structures available for a given chemical series. Answering that question requires a dataset with a deep set of structures in a single campaign rather than broad across targets; Such a dataset would have depositions of ligand intermediates, close analogues, and repeated structures of the same series.

We expand on this work by drawing upon the largest set of publicly available protein-ligand structures for a single target, over 400 crystal structures, generated by the COVID Moonshot Consortium.^46,74^

### The COVID Moonshot provides an unprecedentedly large dataset for cross-docking evaluation

The COVID Moonshot Consortium, a global citizen science effort to combat the SARS-CoV-2 pandemic, started when the Diamond Light Source deposited a crystallographic screen of 71 fragments to the SARS-CoV-2 Main Protease (Mpro) in March 2020.^75^ With the goal of developing these early hits into a potential drug candidate, the Consortium assayed potential hits sourced from worldwide contributors, resulting in potent and developable molecules.^76^ With increased funding, personnel, and scope, the ASAP Discovery AViDD center progressed this program into preclinical studies,^77^ in addition to developing several other discovery programs.^51,52,78^ In doing so, the AViDD center generated an unprecedentedly large, publicly available dataset of protein-ligand crystal structures. This dataset is unique not only for its size, but it also captures a nearly complete drug discovery program from early fragments to optimized leads including progressively more elaborated chemical series developed over time (Figure 2).

**Figure 2:**
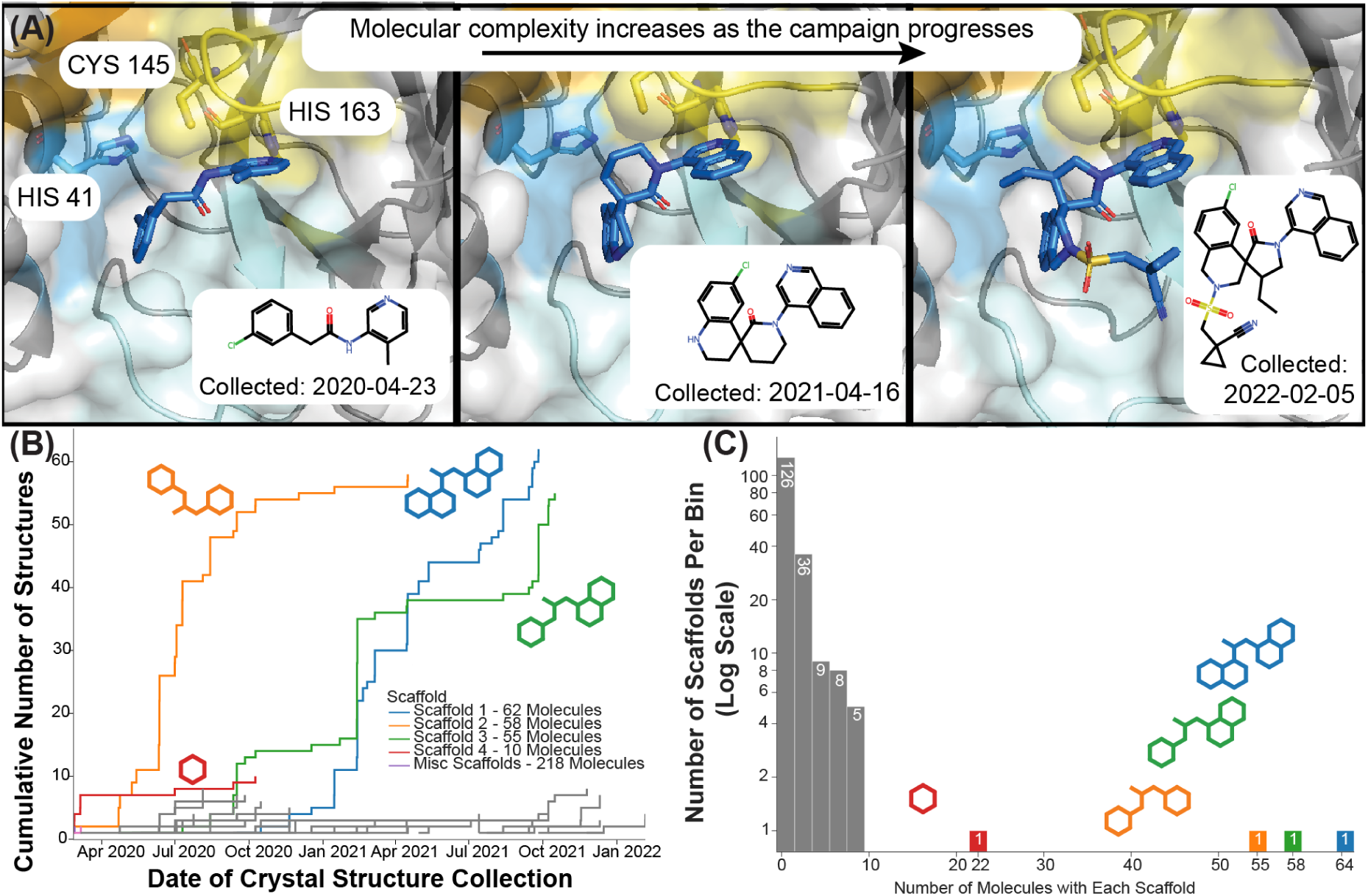
The COVID Moonshot SARS-CoV-2 main protease protein-ligand dataset shows a real-world scenario where the ligands become more elaborated over time. (A) The SARS-CoV-2 main protease binding site in three example crystal structures from the COVID Moonshot showing how the designed ligands increase in degree of elaboration over time. The catalytic dyad of His41 and Cys145, as well as a key P1 residue His163 are shown (sticks). The protein surface is colored by the protease subpocket names S1’ (orange), S1 (yellow), S2 (blue), and S3-5 (cyan). (B) Empirical cumulative distribution of the number of structures per generic Bemis-Murcko scaffolds (corresponding color) over time during the COVID Moonshot campaign. (C) Histogram of the number of molecules per Bemis-Murcko scaffold. Four scaffolds (colored as in B) account for the majority of the dataset, with the remaining scaffolds (gray) representing less than ten molecules each. A significant number of molecules (126) are singlets, not sharing a generic Bemis-Murcko scaffold with any other molecule in the dataset.

Within each scaffold, ligand-to-ligand variation is modest: across the 137 multi-member generic Bemis-Murcko scaffolds, the median maximum heavy-atom count difference is 1, and the largest excursion observed in the entire dataset is 13 heavy atoms (SI Figure S10). For the four most populous scaffolds, median pairwise heavy-atom count differences range from 2 to 4 (SI Figure S11). This relatively conservative chemical excursion within a series supports the use of heavy-atom RMSD as a pose-recapitulation metric within a congeneric set.

The COVID Moonshot dataset represents a promising model system for evaluating reference-based docking performance due to the favorable biophysical properties of the SARS-CoV-2 main protease as an SBDD target.^74^ The SARS-CoV-2 main protease has been the subject of years of inhibitor development studies, with nearly 20 drugs in the market or clinical trials.^77,79,80^ It is a protease with a large, conserved set of protein cleavage sequences, presenting a druggable binding site with many features that have been used to optimize affinity and selectivity.^81^ Relative to other well-studied inhibitor targets, Mpro is readily crystallizable without dramatic modification, amenable to crystal soaking, and the binding site does not undergo any significant conformational changes upon inhibitor binding across different chemical series.^75,82^

Because all 403 protein-ligand structures were obtained via crystal soaking, the energetic cost of large protein rearrangements within the existing lattice limits the dataset to ligands that bind without inducing substantial backbone reorganization. Quantitatively, for the four most populous scaffolds the binding-site heavy-atom RMSD relative to a within-scaffold reference remained below 1 Å and the full-chain RMSD below 2 Å (SI Figure S12). The upper tail of these distributions corresponds almost entirely to comparisons between the monoclinic and orthorhombic crystal forms, rather than to ligand-induced conformational change. This is an important caveat: for targets that exhibit significant induced-fit or cryptic-pocket behavior, more structures per scaffold (or alternative experimental approaches such as cocrystallization) will likely be required than the recommendations in this work suggest.

In this work, we use 403 protein-ligand structures from the COVID Moonshot to quantify how the success rate of accurately predicting ligand poses depends upon the quantity of available structural data. We show that only a small fraction of available structures per ligand series is needed to achieve near-maximal performance. We examine how success rate, defined as pose-prediction within 2 Å of the known pose, varies when only using structures in the order they were collected, replicating a realistic drug discovery campaign. Finally, we explore the consequences of ligand diversity in the structural dataset, identifying how diminishing returns are most evident when collecting additional structures of a particular scaffold. In doing so, we assess which structures are most valuable and identify potential strategies for most effectively leveraging available structural data to support decision making. In summary, we show that scaffold diversity in the structural data is more effective for improving pose predictions than the raw number of structures. This result, demonstrated in a real-world open-science drug discovery campaign, provides for the first time scientific support for a common intuition of medicinal chemists.

## Results and Discussion

### Reference-based docking outperforms standard docking, with the biggest improvement found for ligands docked to similar references

The dataset of crystal structure from the COVID Moonshot was downloaded via Fragalysis on April 1st, 2024.^48^ These 803 crystal structures were filtered to include only non-covalent ligands from the Moonshot discovery project which bind to the active site. This dataset also included duplicates for some structures where the second monomer chain (“chain B”) was aligned to the “canonical” chain A; these were removed to result in a dataset of 576 structures. In the case of duplicate depositions, only the structure and ligand-pose with the earlier collection date were used; this results in a dataset of 414 crystal structures. Other post hoc metrics would use information unavailable during the decision-making point in a live design campaign, and would no longer be a prospective prediction. Of these, 403 were successfully prepared and cross-docked using the OpenEye toolkits as implemented in the *drugforge* software package (see Experimental Section). We refer to the ligand to be posed as the **query ligand**, the crystal structure used in docking as the **reference structure**, and the crystal structure ligand as the **reference ligand**. We used the POSIT algorithm to generate poses of the 403 query ligands against the other 402 reference structures (avoiding the self-docked query-reference pairs) using either the FRED-only or full-POSIT settings, creating two datasets containing 162,006 query-reference ligand pairs with 1 pose each.

For each query ligand, we rank the resulting poses by either the POSIT Probability or by the true RMSD (a positive control) and return the top-ranked pose to evaluate pose prediction success. We define the success rate for pose-prediction as the fraction of query molecules whose top-ranked pose has *≤* 2 Å heavy-atom RMSD to its crystallographic pose, a typical performance metric for pose-prediction studies (Figure 3A).^61,67,70^ RMSD provides a practical performance metric for our *∼*300,000 ligand poses: RMSD is computationally efficient, scales with ligand size, and enables comparison across diverse molecular scaffolds. We note that RMSD as a performance metric is limited as it does not differentiate thermodynamically favorable poses from unfavorable one nor does it consider chemical interactions that favor binding. In other words, a 0.1 Å RMSD pose might contain an implausibly high-energy clash, and a 3 Å RMSD pose might retain all important interactions. RMSD is also blind to the difference between the motions of a flexible substituent exposed to solvent (which may be well-tolerated) or in a constrained pocket. Finally, similar conformational differences can have different RMSD values depending on the total ligand size. For example, given equal motions of a benzene ring in a small fragment and a large ligand, the RMSD of the small fragment will be larger. As such, while RMSD is an imperfect metric that requires careful consideration during interpretation, it remains a tractable, scalable, and efficient metric for our diverse library of chemical moieties.

**Figure 3:**
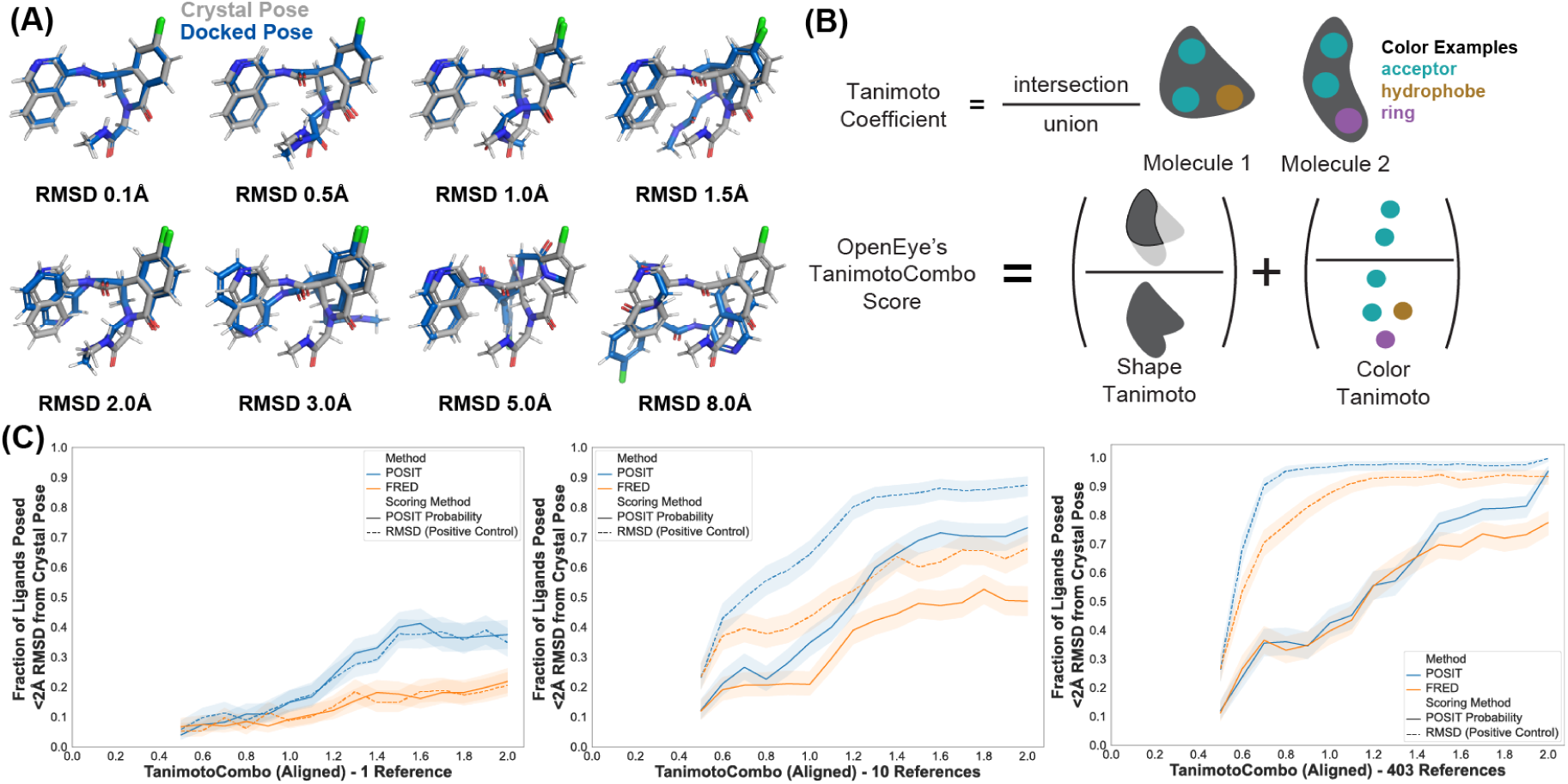
Pose prediction accuracy improves with increasing ligand similarity to the reference structure. (A) Examples of docked ligand poses (blue) with increasing RMSD to the crystallographic pose (grey) for an example ligand (MAT-POS-a54ce14d-2). (B) The TanimotoCombo Score used by OpenEye’s POSIT docking algorithm combines the Shape and Color Tanimoto Coefficients to provide a scaffold-independent, spatially-aware measure of chemical similarity (see Methods). (C) Pose prediction success is plotted as a function of the similarity of the query ligand to the reference ligand, comparing two docking methods: POSIT (blue) and FRED (orange), and two scoring methods: the RMSD as a positive control (dashed line) and the POSIT Probability (solid line). The success rate is shown for three examples, in which 1, 10, or all available structures (403) are available to use. For each query ligand, the structures are first filtered by similarity, and then the number of references indicated are randomly sampled from the remaining structures by bootstrapping 1000 times. Although POSIT outperforms FRED for similar molecules, this relative improvement decreases when more structures are used. Sample sizes per TanimotoCombo bin are reported in the histogram beneath each panel. The narrowing of the bootstrap confidence intervals at high TanimotoCombo reflects both the increased bin population and the saturation of the success rate near 1; the bands should therefore be interpreted as joint bin-count and binomial-variance intervals, not as evidence of equal sample size across bins.

We measure the success rate as a function of the aligned TanimotoCombo similarity for both the FRED and POSIT docked poses (Figure 3C) and find that the performance of reference-based docking (POSIT) outperforms standard docking (FRED) as the molecules become more similar.^71^ We first reproduce previous findings that show that POSIT out-performs other docking methods when docking to similar reference ligands (Figure 3C). ^67,71^ While FRED relies solely on protein-ligand interactions, POSIT incorporates Tanimoto-Combo scoring to align the query ligand with the reference (crystal structure) ligand. We would expect the POSIT-posed success rate to increase more than the FRED-posed success rate as a function of increasing query-to-reference ligand similarity since the protein structures in this dataset are quite similar to each other. This expectation holds when using a single reference structure (Figure 3C, left panel), where the improvement in the success rate as a function of the ligand similarity is larger for full POSIT (blue, 0.05 to 0.35) than for FRED (orange, 0.05 to 0.15). However, when all possible references are included, the performance is equivalent until the similarity is *>*0.8 (Figure 3C, right panel). This suggests that FRED’s performance approaches that of POSIT when FRED is given enough opportunities to generate poses via having additional reference structures.

We examine a few ways of measuring chemical similarity, including the TanimotoCombo score from OpenEye (Figure 3B),^67^ the maximum common-substructure (MCS) (SI Figure S2A),^83^ and the extended-connectivity fingerprint (ECFP)^84^ (SI Figure S2B). The aligned TanimotoCombo and MCS Tanimoto coefficients (Figure 3B, Methods) describe the dataset as being less diverse, while the ECFP4 fingerprint and ECFP10 fingerprint describe the dataset as being more diverse (SI Figure S2C). The ECFP Tanimoto measures chemical similarity more in terms of the presence of functional groups and local motifs, while the MCS Tanimoto is more sensitive to changes in the core scaffold. Since a majority of core scaffold diversity stems from variations of the decorating groups, the fingerprints report greater ligand diversity than MCS Tanimoto. We use the TanimotoCombo as our measurement of chemical similarity, as it provides the widest range of ligand similarity scores, avoids the idiosyncratic behavior and feature choices of the MCS, and takes advantage of available 3-dimensional information.

The discrepancy between the POSIT Probability (Figure 3C, solid lines) and the true RMSD (Figure 3C, dashed lines) increases when more references are used. When 10 references are available, the maximum difference between the POSIT Probability (0.25) and the true RMSD (0.6) is at a TanimotoCombo similarity of 0.8, with a difference of 0.35. This difference grows to 0.6 when all 403 reference structures are used. This suggests an improved scoring method for query-reference pairs with a TanimotoCombo of *<*0.8 would be the most impactful. We also performed this comparison with the Tanimoto of the ECFP4 2048 bit fingerprint as implemented in RDKit (SI Figure S2D). Upon analyzing the success rate as before, we observe that the success rate plateaus after similarities of *∼*0.4, consistent with prior results (SI Figure S2C). There are few query-reference pairs with an ECFP4 similarity of *>*0.4. Thus, we suggest against using ECFP4 as a similarity metric for analyzing single-target datasets, as it conflates the chemical diversity represented by the dataset.

Across the 414-structure dataset, ligand diversity is captured at four complementary levels. At the scaffold level, we identify 137 distinct generic Bemis-Murcko scaffolds (Figure 2C), of which the four most populous account for the majority of the ligands (Figure 2A,B). Within each scaffold, chemical excursion is modest — the median maximum heavy-atom count difference within a scaffold is 1, and even the worst-case within-scaffold pair differs by only 13 heavy atoms (SI Figure S10). Pairwise chemical similarity, measured by aligned TanimotoCombo, MCS atom count, and ECFP fingerprints, places the dataset in a moderate-similarity regime (SI Figure S2C). Finally, the dataset spans a correspondingly broad range of protein-ligand interaction fingerprints (SI Figure S14), so that the cross-docking analysis explores genuinely distinct interaction patterns rather than minor variations on a single binding mode.

It is worth noting that success rate includes structures drawn from the whole dataset, which does not reflect the prospective nature of a drug discovery campaign. Rather, the dataset of a successful drug discovery campaign will inherently converge into a smaller subset of molecule designs. As such, prospective assessments must consider the time-evolution of ligand design. To appropriately estimate the prospective performance of reference-based docking, we consider the time-evolution of the Moonshot dataset by comparing a random selection of the structures to a selection ordered by the date of crystal structure collection.

### Pose prediction success has diminishing returns with an increasing number of available crystal structures

To assess how increasing the number of crystal structures improves pose prediction performance, we compare two splits in which structures were ordered either randomly (Random Split) or chronologically by collection date (Temporal Split) then incrementally added to the pool of available structures for each reported success rate (Figure 4A). The Temporal Split provides a simulation of a prospective campaign: at each point along the horizontal axis, only the structures actually solved by that date are available for docking. The time-split establishes when additional structures stop changing the decision a project team would make. Findings from a temporal split do not capture improvements in crystallographic technology improvements in crystal-structure determination would require comparing depositions across many targets and years. This would involve varying the target, preparation, and chemical series, which differs from our campaign but have been used in previous efforts and benchmarks to assess docking performance.^72^ Performance is better for the randomly shuffled dataset (Figure 4B) since randomly shuffling structures increases the probability of the query and reference molecules having a high similarity.

**Figure 4:**
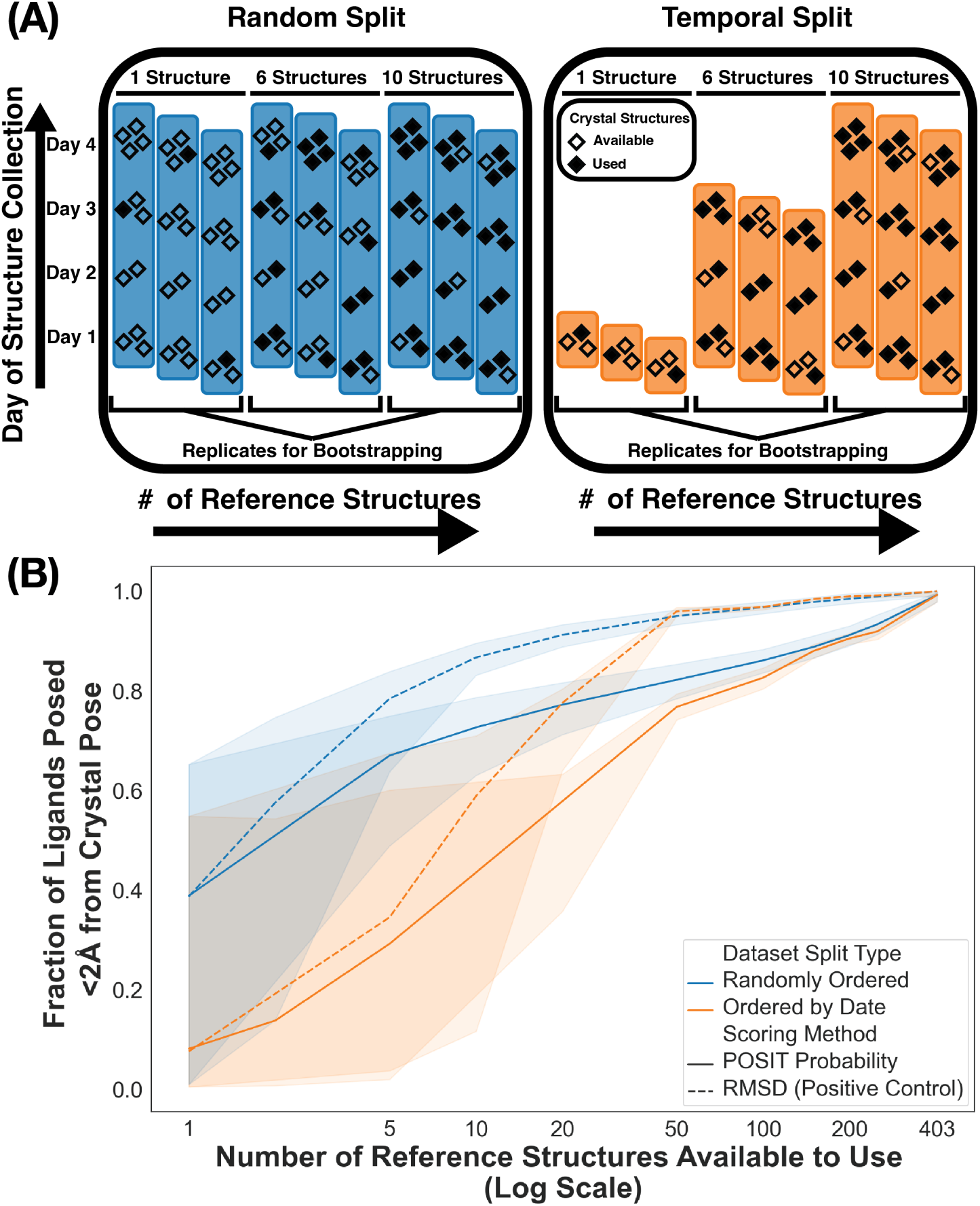
Pose prediction is more challenging when limited to the first crystal structures collected during the campaign. (A) Schematic representation of the two dataset splits. For the Random Split (blue, left), reference structures are randomly selected from the entire dataset (highlighted blue regions), whereas for the Temporal Split (orange, right), possible reference structures are selected chronologically by increasing the last date of crystal structure deposition, with same-day structures randomized. For both splits, the confidence interval is reported by bootstrapping over 1000 samples. (B) The success rate (Fraction of Ligands Posed *≤* 2 Å from the Crystal Pose) for both dataset splits are reported as a function of the number of reference structures available for pose prediction. Success rate is shown for both the Random Split (blue) and Temporal Split (orange) and for poses ranked either by the RMSD to the crystal pose (solid line) and the POSIT Probability (dotted line). Bootstrap intervals (shaded bands) act as a proxy for the number of reference structures available at each count; their width at small reference counts reflects the dependence of the outcome on which particular structures have been solved, and their contraction with increasing count is the effect under study.

With the first 50 crystal structures collected during the COVID Moonshot Initiative (in the Temporal split), 75% of the ligands can be posed within 2 Å of their crystallographic pose using the POSIT docking algorithm (Figure 4B, orange solid line). As before, we compare ranking the predicted poses for each ligand by the POSIT probability to ranking them by the RMSD as a positive control. For the RMSD-ranked poses, the difference in success rate between the two structure-splits is *<*5% after *∼*20 structures, but it takes *∼*120 structures for the POSIT-ranked poses to match the performance of the RMSD-ranked poses.

There is a large distribution of success rates for the Temporal Split when using *<*10 structures (Figure 4B, orange lines). Some initial structures result in success rate *<*10% independent of ranking by RMSD or POSIT probability, suggesting that initial failures are due to posing rather than scoring.

Although only an eighth of the dataset is needed to get above a 70% success rate in the Temporal Split, some ligands are not posed correctly until the entire dataset of structures is used. To further examine the causes of these failed pose-prediction tasks, we first explored how success rates depend on the scaffold used.

### Few structures per scaffold are needed to correctly pose a candidate molecule

We hypothesize that the main source of diminishing returns stems from adding structures with the same generic Bemis-Murcko scaffold. That is, for an individual scaffold, additional structures collected do not significantly improve success rate and rather offer costlier diminishing returns. For robust statistics and representation, we limit this analysis to scaffolds with at least 10 molecules represented in this dataset, which include the top four most common scaffolds (Figure 2C). We calculated the success rates for these four scaffolds docked only to structures with the same scaffold. Even with only 1 structure, the success rates were higher for the first three scaffolds than for the full split (Figure 5A-C). For these three scaffolds, we observe minimal improvement in success rate after the second structure is added suggesting that additional corresponding structures of a given scaffold are not needed to achieve a maximal success rate. On the other hand, scaffold 4, constituting single ring fragments, is an outlier. Each added structure provides a significant improvement in the fraction of ligands posed within 2 Å of the crystal pose (Figure 5 D). This is consistent with previously observed challenges in fragment pose prediction tasks.^85–87^

**Figure 5:**
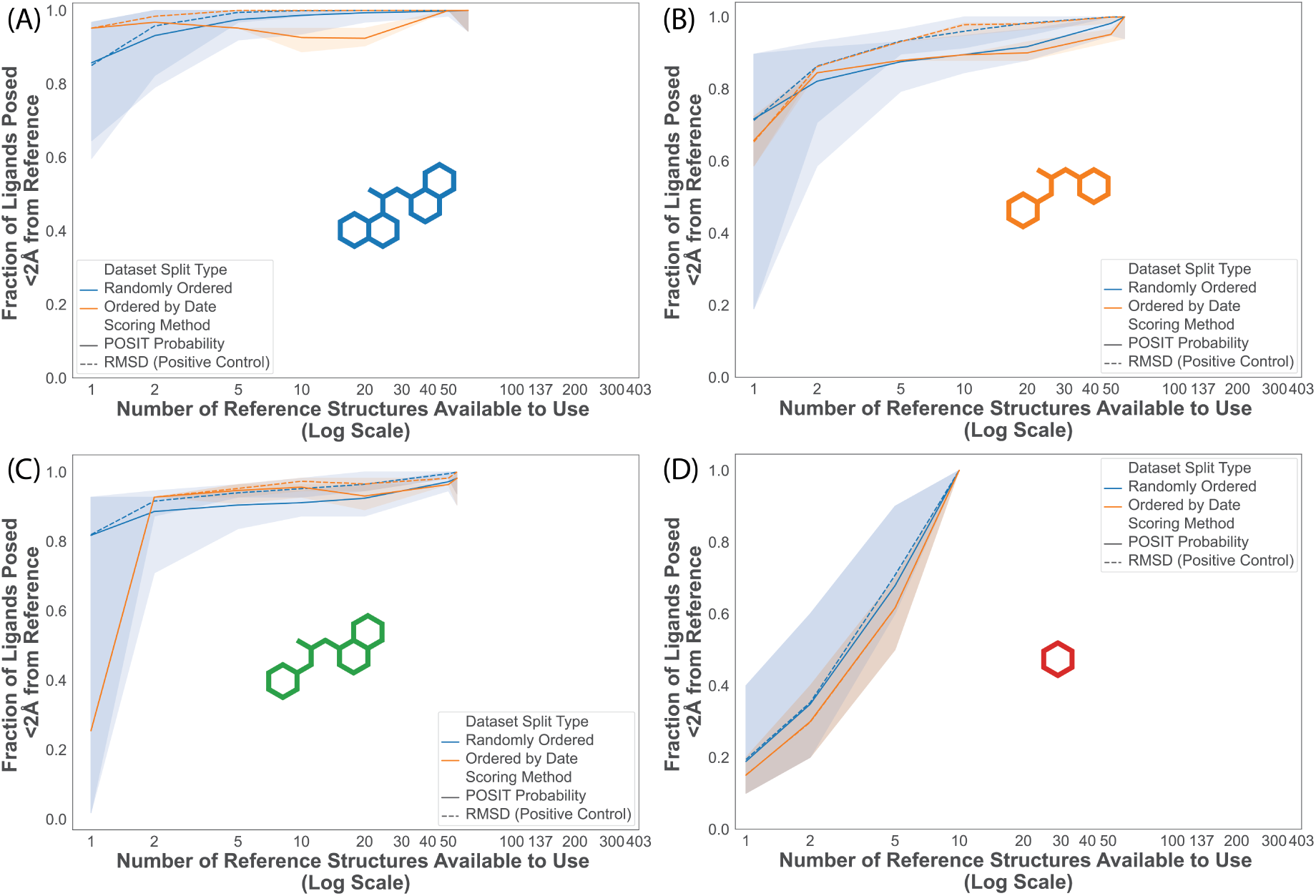
Pose prediction success is higher when the reference and query have the same scaffold, unless the scaffold is a single-ring fragment. The success rate (Fraction of Ligands Posed *≤* 2 Å from the Crystal Pose) of scaffolds 1-4 (A-D, respectively) are reported as a function of the number of reference structures available for pose prediction. The total number of structures is different for each scaffold but they are plotted on the same scale as Figure 4 for clarity. Success rate is shown for both the Random Split (blue) and Temporal Split (orange) and for poses ranked either by the RMSD to the crystal pose (solid line) and the POSIT Probability (dotted line).

We assess whether pose prediction performance can be maximized in sparse data regimes by limiting the number of structures collected for any ligand scaffold. We analyze this strategy (Figure 6, green lines) in comparison to the Temporal Split (Figure 6, orange lines). Below 50 structures, the efficiency between the different strategies remains similar, while we do see a small improvement in efficiency beyond 50 structures. However, the success rate has already reached *∼*80% when over 50 structures are present leaving only minimal room for improvement. To examine whether additional structures per scaffold improve the performance, we simulate collecting an increasing number of structures per scaffold, from 1 to 20, evaluating efficiency as a function of the number of available structures. We do not see any significant improvement in efficiency as the number of structures per scaffold increases (SI Figure S8).

**Figure 6:**
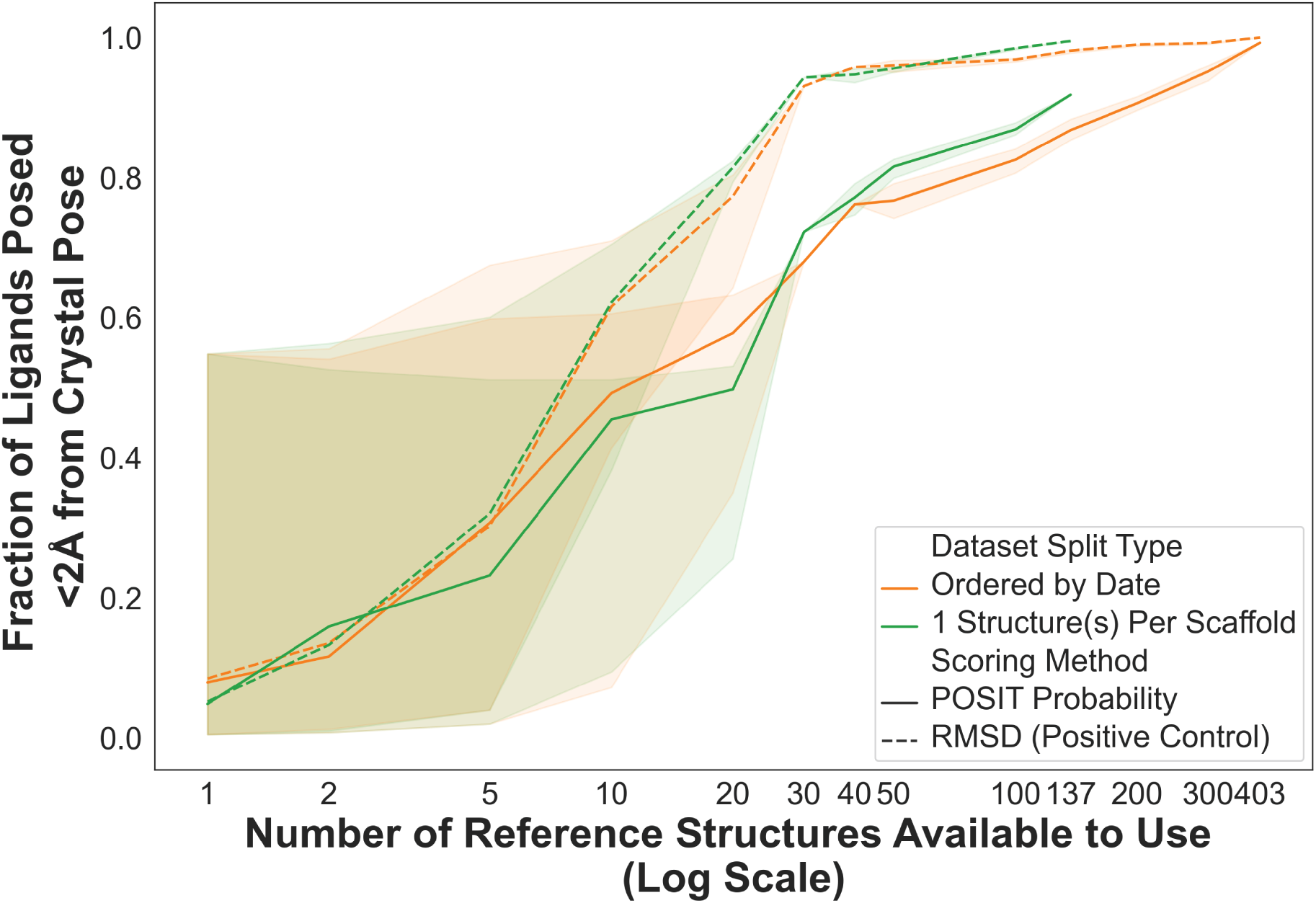
Collecting only 1 structure per scaffold shows a minor efficiency improvement from the temporal split beyond 50 structures. Pose success rate (fraction of ligands posed *≤* 2 Å from the crystal pose) is reported as a function of the number of reference structures available for pose prediction. Success rate is shown for the previous Temporal Split (orange) and when a single structure per scaffold is made available (green) for poses ranked either by the RMSD to the crystal pose (solid line) and the POSIT Probability (dotted line). The structures for the One Structure Per Scaffold Split are made available in order of their collection date, as in the Temporal Split.

To further probe whether any particular scaffolds were more challenging than others, we analyze each crosswise query-reference pair for the top 20 scaffolds, ranked either by RMSD (SI Figure S4A) or by POSIT Probability (SI Figure S4B). By averaging the results we see that for all the scaffolds, predicting their poses was more successful than using their crystal structures as the reference (SI Figure S5).

As observed previously, the single ring fragment (scaffold 4) remains challenging both as query and as reference (SI Figure S7, top left). Scaffold 6 (SI Figure S7, top row, middle) was also challenging for both, likely due to the extended linker between the central amide bond and the single ring system, giving it increased flexibility. Scaffold 8 (SI Figure S7, top right) is unusual in this dataset as it contains a *β*-lactam ring, which may have contributed to its challenges for pose prediction. Scaffold 14 (SI Figure S7, bottom left) is also notable as it contains one of the few examples of a branching point with a tertiary carbon. This moiety was also explored with scaffold 33 (SI Figure S7, bottom right), which also performed poorly.

Recapitulating the geometric pose within 2 Å RMSD does not guarantee that the underlying protein-ligand interactions are preserved. We therefore re-evaluated the analyses of Figures 3C, 4B, 5, 6, and 7 using a PLIF Recall of 1 as the success criterion (SI Figure S14). We define the PLIF Recall for a docked pose as the fraction of protein-ligand interaction fingerprint (PLIF) bits present in the reference crystal pose that are also recovered in the docked pose, where PLIFs are computed using a standard interaction definition (hydrogen bonds, hydrophobic contacts, aromatic stacking, ionic interactions, and halogen bonds within their respective distance and angle cutoffs); a PLIF Recall of 1 indicates that every interaction made by the crystallographic ligand is recovered in the docked pose. The qualitative trends are unchanged: success rates increase with reference-ligand similarity and with the number of available reference structures, and POSIT continues to outperform FRED at high similarity. Absolute success rates are uniformly lower under the stricter PLIF criterion, but the diminishing-returns plateau as a function of the number of available structures is preserved, supporting our central claim that scaffold diversity, rather than within-scaffold depth, drives most of the gains.

In conclusion, we identify a point of diminishing returns by limiting our dataset to 1-2 structures per generic Bemis-Murcko scaffold. However, for a project such as the COVID Moonshot which generated *>*100 generic Bemis-Murcko scaffolds, this would still require collecting *>*100 structures. To avoid having to collect *>*100 structures, we wanted to find a set of 5-10 structures with a high success rate for the whole dataset. We show that a small number of same-scaffold structures brings the three elaborated scaffolds close to their plateau, exceeding 95% success for scaffold 1 and reaching 88–90% for scaffolds 2 and 3 at five structures (Figure 5A-C, SI Table S1). Therefore, we next evaluated how ligands from the remaining scaffolds perform when docked to these structures (SI Figure S18). With a success rate of 60% at 10 structures, collecting several structures of the top three scaffolds is a reasonable strategy for using a minimal number of structures to achieve the best possible performance. However, it remains unclear whether docking failures are primarily due to failures in pose sampling, or in failures to rank multiple poses.

### Keeping up to 50 poses from POSIT shows a minimal improvement in success rate even with a perfect ranking

So far we have only analyzed pose prediction results where a single pose is returned by the docking algorithm. Retrieving multiple poses for each protein-ligand complex hurts scalability and risks a computational pipeline becoming prohibitively expensive for a real-world drug discovery campaign. To test whether the failures stem from poor initial pose generation or inaccurate POSIT probability scoring, we had POSIT return up to 50 deduplicated poses for each query–reference combination (see Experimental Section). The success rate is plotted, as before, as a function of the number of reference structures available to use for either the Random or Temporal Split (Figure 7). Since the POSIT Probability is used to rank poses by default, we report the positive control (RMSD-ranked) success rate. As before, the Random Split has a higher success rate than the Temporal Split, and reference-based docking (POSIT) outperforms protein-only docking (FRED). Increasing the number of poses returned by the docking algorithm from 1 to 50 improves the success rate, with the greatest improvement for both methods occurring when 5–20 structures are made available. FRED shows a 20% improvement and POSIT has a 10% improvement in pose prediction success rate. This effect is smaller than that seen for adding more structures, matching results shown in previous kinase cross-docking benchmarks.^71^ Our results highlight the importance of pose-sampling; with current pose sampling strategies, a perfectly predictive scoring function would only improve success rate by 10-20%. The poor success rate observed at the leftmost portion of the Temporal Split is not a failure of POSIT per se, but a reflection of the chemistry available at that stage of the campaign. The earliest 5–20 crystal structures are dominated by small single-ring fragments (scaffold 4) with fewer than *∼*15 heavy atoms; pose prediction for fragments of this size is intrinsically difficult regardless of method. Once the campaign progresses to more elaborated scaffolds (*∼*10 additional structures), Temporal-Split performance converges with Random-Split performance (SI Figure S13), confirming that the temporal effect is largely driven by fragment size and scaffold availability rather than by intrinsic time dependence.

**Figure 7:**
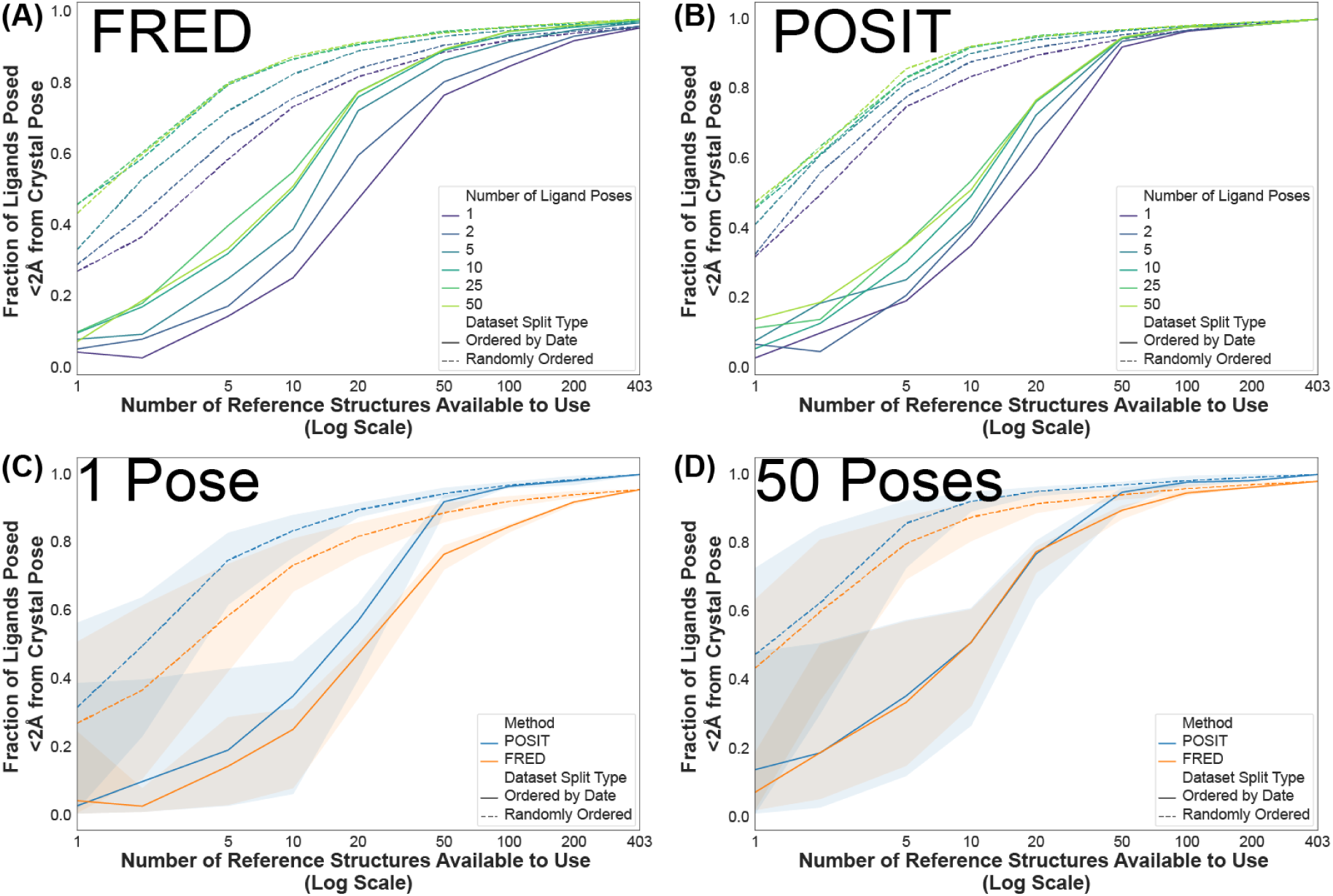
The lowest RMSD pose generated by the POSIT algorithm may not be ranked best by the POSIT probability. (A) and (B) The success rate (fraction of ligands posed *≤* 2 Å from crystal pose) is reported as a function of the number of reference structures available to generate up to 50 ligand poses using either FRED (A) or POSIT (B) and ranked by RMSD. The number of poses generated range from 1 (purple) to 50 (lime green) and structures used were using either a Random Split(dashed line) or a Temporal Split (solid line), as in Figure 4. (C) and (D) The same data is plotted for 1 pose (C) and 50 poses (D) for FRED (orange) and POSIT (blue) generated poses for the random (dashed line) or ordered-by-date (solid line) structures. These results show that ranking up to 50 poses by even a perfect scoring function (RMSD, the same function we use to measure success) only increases the success rate by 10%, and with more than 50 structures there is no significant improvement to generating more poses. The improvement in the success rate is more significant for the FRED-posed structures (20% → 50%) than the POSIT-posed structures (30% → 50%).

We deliberately frame this study around pose prediction rather than affinity ranking. Docking scores typically correlate only weakly with experimental binding affinity at the single-compound level and are most useful as coarse filters applied to libraries of more than 10^6^ molecules; with approximately 400 ligands the within-scaffold variance of POSIT and ChemGauss4 scores is too large to support reliable affinity-ranking conclusions. Achieving robust statistical power to evaluate the impact of pose-prediction accuracy on docking-score-based affinity ranking would require at least an order of magnitude more protein-ligand structures of a single target than is currently available in the public domain. We therefore restrict the present analysis to pose prediction, where the available sample size is sufficient.

Ligand B-factor here denotes the mean isotropic atomic displacement parameter over the ligand heavy atoms, in units of Å^2^.^88^ B-factors are unbounded above and distinct from other normalized real-space fit measures such as Real-Space Correlation Coefficient (RSCC) and Real-Space R-value (RSR) that also appear in PDB validation reports.^89^ Larger B-factor values correspond to greater positional uncertainty of the modelled ligand atoms.

The Moonshot structures were solved in two crystal forms, denoted in the deposition as the x-series and the p-series, corresponding to two soaking systems used at different stages of the campaign. Crystal form describes the crystallization and soaking conditions rather than the quality of the resulting model and is not a validation metric.

To interrogate the effect of crystal structure quality on pose prediction success, we have plotted the average success rate per reference (across all query ligands) as a function of the ligand B-factor (SI Figure S16), as a function of the R-factor pooled across crystal forms (SI Figure S20), and the pairwise relationships between the standard quality metrics (SI Figure S19). The per-reference success rate was more significantly impacted by which crystal form it was in than by the ligand B-factor, a trend that held for other quality metrics such as resolution and R-factor. In fact, despite the worse crystal quality metrics for the orthorhombic (p-series) structures (SI Figure S17), these structures generally outperform the monoclinic (x-series) structures, likely due to the time-dependent chemical similarity trends discussed elsewhere in this article. We observe that crystal quality does not improve monotonically over the campaign, the later orthorhombic structures carry higher ligand B-factors than the earlier monoclinic ones. As such, pose prediction success cannot be attributed to improving model quality. We acknowledge that this conclusion may not extend to datasets with broader quality variability, particularly those including structures with poorly resolved ligand density.

### Limitations and scope of applicability

Our quantitative recommendations are derived from a single protein target and should be read as a lower bound rather than a universal threshold. All 403 structures of the SARS-CoV-2 main protease appear to have a rigid binding site: heavy-atom RMSD to the within-scaffold reference remains below 1 Å in the site and below 2 Å across the entire chain (SI Figure S12). Targets that undergo induced-fit rearrangement or cryptic-pocket opening will require more structures per series than we report; previous kinase-specific cross-docking benchmarks show systematically lower success at matched structure counts.^71^ Where soaking is not viable and co-crystallization is needed, the cost of each structure also rises, shifting the resource trade-off that motivates this analysis.

Our benchmarks were measured using a single docking platform and absolute success rates are engine-specific. POSIT and FRED share preparation, scoring, and file-handling software pipelines, isolating template availability as the only variable that differs across these comparisons. Using an engine implemented using a different software stack risks mixing protocol differences alongside measurement effects. A second independent template-based method, such as FitDock^73^ and Surflex-Dock,^72^ would establish whether the plateau is a property of the underlying data or of POSIT’s shape-and-colour matching; This would be a valuable extension of our current work to establish generalizability of our observed trends. That previous efforts using FitDock and Surflex-Dock report a template-driven advantage over template-free docking suggests our observations are not specific to POSIT.

Our recommendations apply to congeneric series as they were realized in this campaign, and so generalizability should be carefully considered. One generic scaffold can in principle span an enormous range of chemical substitution. In our dataset, across the 137 multi-member scaffolds, the median maximum within-scaffold heavy-atom difference is one atom, and the largest deviation across a scaffold series for any pair of compounds anywhere in the set is 13 atoms (SI Figures S10 and S11). Design campaigns that diversify rapidly within a single scaffold might thus expect to need more crystal structures to template against than we find in our instance.

Our strongest observed findings occur on the ligand scaffold-series that advanced the furthest and may not apply to other scaffold-series. Scaffold 1 is both the most populous series in the Moonshot dataset and reaches greater than 95% pose prediction success with five same-scaffold structures (SI Table S1). That the scaffold 1 series was selected under a real resource-allocation constraint provides useful guiding principles for future prospective campaigns; the number is conditioned on the properties that made the series worth advancing. The two other elaborated series reach 88% and 90% at five structures and require roughly an order of magnitude more structures to exceed 95% success rate. A project working on a scaffold-series may find it useful to treat five-structures per scaffold as a point of diminishing returns in resource cost for a virtual screening campaign rather than the point at which accuracy is assured.

The Temporal Split assumes that deposition order tracks the design-make-test-analyse progression of the campaign, an assumption specific to the COVID Moonshot. Structures were solved as compounds were prioritized and synthesized, so a later structure corresponds by construction to a later design decision, and the published campaign record supports this.^74^ Campaigns that solve structures in batches, or in which crystallography lags synthesis by months, will map structures onto campaign stage differently, and the split should be read as a simulation of accumulating evidence rather than as a calendar. Furthermore, our temporal split cannot capture or assess performance dependence through improvements in crystal-structure determination and crystal model quality; such an effort would require a cross-target design spanning many campaigns and could not hold the target, the protocol, or the chemical series constant. Additionally, our finding that success does not degrade with crystallographic quality holds only across the range this dataset spans. Resolution runs from 1.2 to 2.5 Å, R-free from 0.19 to 0.33, and mean ligand B-factors from roughly 15 to 80 Å^2^, and these metrics are strongly intercorrelated (Spearman *ρ* = 0.60–0.91; SI Figure S19). Within our dataset resolution range, per-reference success rates do not decrease as crystal structure quality worsens (SI Figure S20). Crystal form and campaign stage are also confounded: the monoclinic x-series structures were collected early and carry the smaller fragments, while the orthorhombic p-series structures were collected later, carry the elaborated ligands, and show higher ligand B-factors. Therefore, we cannot attribute differences in their performance to crystal structure quality, form, or ligand chemistry. Datasets containing poorly resolved ligands fall outside this range and the result should not be extended to them.

Lastly, we emphasize that our study addresses pose prediction under a single geometric success criterion. Docking scores correlate weakly with binding affinity at the single-compound level, and our sample of 400 ligands may be too few to support affinity-ranking conclusions. Such questions likely require orders of magnitude more single-target structures than exist in the public domain.

## Conclusion

In this work, we use a publicly visible drug-discovery-campaign—the COVID Moonshot— to retrospectively evaluate the effect of the number of available crystal structures on pose prediction performance. We find that only a few structures of any scaffold-bound target are needed to successfully predict the poses of future candidate molecules later in the campaign. Matching chemical intuition, this performance is dependent on the similarity of the reference to query ligand. While reference-based docking (POSIT) significantly outperforms protein-only docking (FRED) when only a few structures are available, the difference becomes less significant as more structures are included. Although the fraction of correctly predicted poses increases with the number of structures used, this improvement results in diminishing returns such that only a fraction of available structures are needed to approach maximal performance. In particular, predicting the pose of a molecule with a given generic Bemis-Murcko scaffold is most successful when docking to a structure with that same scaffold, supporting the commonly held medicinal chemistry intuition that scaffold diversity is more important than the raw number of structures. The POSIT Probability is moderately predictive of pose prediction success, with failures stemming from poor pose generation (Figure 4B). In the Temporal Split, the RMSD-ranked success rate is only ever about 20% better than the POSIT-Probability ranked success rate. Our pipeline-driven analysis highlights that while few structures are needed to pose 70% of ligands accurately, a poor match between the reference ligand and the query ligand results in the majority of failure modes. It is worth noting that the success rate required will likely depend on the specific target and the downstream computational method being used.

In conjunction with our retrospective analysis, we provided the baseline pose-prediction model for a prospective blind-challenge for SARS-COV-2 and MERS-CoV pose prediction, in collaboration between Polaris, OpenADMET, and ASAP Discovery.^90^ Although our docking workflow was the best performing non-co-folding model, the top three best performing models used co-folding.^90^ Similarly, co-folding models have recently been shown to be competitive with docking models for preparing structures for free energy calculations although at much greater computational cost.^91^

Recent co-folding methods (AlphaFold3, Boltz, Chai, and successors) outperform classical template-based docking on certain prospective benchmarks, particularly for ligand series that are well represented in their training data. For ligand series outside the training distribution, however, co-folding methods can produce confidently wrong poses; recent work documents specific failure modes for genuinely novel scaffolds.^41,92^ Classical template-based docking remains advantageous in two regimes: (i) when high-quality crystal structures of similar reference ligands exist, where the inductive bias provided by the template is exactly what we have shown improves pose prediction; and (ii) when uncertainty quantification is required and the explicit dependence on a chosen reference makes the failure mode interpretable. The recommendations of the present work—prioritize scaffold diversity in the experimental dataset—are largely orthogonal to the choice of downstream method and apply to co-folding workflows as well.

The widening gap between the POSIT-Probability-ranked success rate and the RMSD-ranked oracle as the number of available reference structures increases (Figure 4B) is informative. Adding more reference structures primarily helps the sampling step—a correct pose is generated more often—while the ranking step (selecting the correct pose from a growing pool of plausible alternatives) is not equivalently improved. This decomposition supports our broader observation that pose generation, rather than pose scoring, is the dominant bottleneck in the regimes explored here, and motivates future work on ranking methods that scale better with reference-pool size.

Future extensions of our work would evaluate the impact of pose prediction performance on the accuracy of alchemical free energy estimation.^57^ Additionally, future work will explore the use of protein-ligand interaction fingerprints to evaluate pose prediction performance.^92,93^

In the interest of providing practical guidance to drug hunters in light of our results, we have several recommendations. First, we have seen that pose prediction for small fragments is not likely to be successful, so experimental determination of early hits is highly valuable. Second, since additional structures improve pose prediction the most for new generic Bemis-Murcko scaffolds, we recommend solving new crystal structures each time a new scaffold is proposed. Finally, we have seen only minimal benefit from the addition of further structures after 5–10 for a given scaffold, so we recommend instead prioritizing the collection of new scaffolds. However, we recognize that these recommendations may not hold for protein targets where crystal soaking is not viable due to high flexibility in the binding site.

Future versions of our analysis would benefit from large datasets of protein-ligand structures for more challenging targets. While large amounts of structure datasets are more accessible to academic groups than been before, many proprietary protein-ligand structural datasets are maintained by pharmaceutical companies.^94^ Thus, a vast majority of single-target protein-ligand datasets, and the downstream discoveries they enable, are locked away as closed-source repositories. However, there is ample evidence that availability of protein-ligand structural datasets enables discovery; compounds from the COVID Moonshot were the foundation for other groups to develop their own SARS-CoV-2 Mpro inhibitors.^95^ Realizing this, several of these companies have joined together to use “federated” machine learning on their private datasets, enabling them to benefit from this vast source of data without needing to disclose it.^96,97^ Academic groups such as the Structural Genomics Consortium^98–100^ and the newly minted OpenBind Consortium^47,101^ are working towards generating large datasets of protein-ligand structures for clinical targets in an open-source, open-science manner. We hope this contribution to the field will help guide these groups to collect the structures which provide the most value for structure-based drug discovery.

## Methods Section

### SARS-CoV-2 Protein-Ligand COVID Moonshot Dataset

The dataset of 803 protein-ligand crystal structures collected by the COVID Moonshot Consortium were downloaded from the Fragalysis website using their API on 2024/04/01 using scripts from the *asapdiscovery* software repository. Of these, just the 366 X-series and 265 P-series structures were further filtered by selecting only the * 0A structures, which contain all crystal structures in which the ligand is bound to Chain A of the SARS-COV-2 Main Protease. To further refine this, only the structures in which the ligand is bound to the active site of the protein were selected, yielding 543 protein-ligand crystal structures.

#### Structure Preparation with Spruce TK

The structures were prepared using the OpenEye Toolkit (version 2022.1.1) according to the following steps: 1) building in sidechains, loops, missing residues, and hydrogens, and 3) protonating any protonatable residues. This preparation was performed using code from the asapdiscovery toolkit.

#### Crystal structure quality metrics

Resolution, R-factor, R-free, and per-atom isotropic displacement parameters were retrieved for each deposited entry from the RCSB PDB REST API and the corresponding wwPDB validation reports.^89,102^ Ligand B-factor is reported throughout as the arithmetic mean of the isotropic displacement parameters of the ligand heavy atoms, in Å^2^; no radius-based selection of surrounding protein atoms was applied, and no normalization was performed. Retrieval and aggregation were carried out with scripts available in the repository listed in the Data Availability Statement.

### OpenEye Docking

POSIT was selected for this study because it is a template-based docking method that explicitly uses reference crystal structures of similar ligands during pose generation; it is therefore directly suited to the scientific question of how reference-structure availability influences pose prediction. FRED was used as the template-free baseline because it shares POSIT’s structure preparation, scoring, and file-handling pipeline, isolating the use of template information as the only altered variable. Engines built on independent software stacks, whether template-free (e.g., AutoDock) or template-based (e.g., FitDock,^73^ Surflex-Dock^72^), were not included, since differences in preparation and scoring would convolve with effects under study. Extending this analysis across docking platforms is an important direction for future study (see Limitations). OpenEye’s POSIT algorithm was used to dock the structures, with helper code from the asapdiscovery toolkit. This is a rigid-body docking algorithm that uses either FRED, HYBRID, or SHAPEFIT algorithms from OpenEye depending on how similar the query ligand is to the reference ligand. Although multiple structures can be passed to POSIT to enable automated choice of optimal reference structure, for the cross-docking analysis, each pairwise combination of ligand and protein were docked separately (excluding the self-docked query-reference pairs), resulting in 162,006 (403 *×* 402) potential docked structures. The ligands were passed to this docking function as 2D SDF files to avoid biasing the docking results with the crystalized conformation of the ligand. For the multiple-pose analysis, 50 poses for each protein-ligand combination were requested from POSIT, however in many cases, the algorithm will return less than the requested number of poses, with some being duplicates. To account for this, all poses were ranked according to POSIT probability, and any pose with an RMSD *≤* 2 Å to any other, higher ranking pose was removed, resulting in a set of deduplicated poses. This is motivated by the fact that since RMSD follows the triangle inequality, if pose A is *≤* 2 Å to the crystal structure pose, and *≤* 2 Å to pose B, then pose B is also *≤* 2 Å to the crystal structure.

### Chemical Similarity Analysis

#### Maximum Common Substructure (MCS)

The Maximum Common Substructure was calculated for all ligand pairs using the OpenEye python toolkit using default settings. The MCS Tanimoto Coefficient was calculated as the number of atoms in the MCS divided by the total number of atoms in both ligands minus the number of atoms in the MCS.

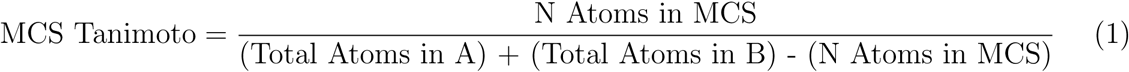

#### Extended-Connectivity Fingerprint (ECFP)

The ECFP was calculated for all ligand pairs using the OpenEye python toolkit. The results shown in this paper were calculated with a radius of 2 (ECFP4) or 5 (ECFP10) and a bit size of 2048. The Tanimoto Coefficient was calculated with the OpenEye function OETanimoto.

#### TanimotoCombo Score

The TanimotoCombo score was calculated for all ligand pairs using the OpenEye python toolkit. The TanimotoCombo Score used by OpenEye’s POSIT docking algorithm combines the Shape and Color Tanimoto Coefficients, which provides a scaffold-independent, spatially-aware measure of chemical similarity. The shape is defined by the combined Gaussians used to represent the ligand atoms, and the color is defined by the Mills Dean Implicit Force Field and includes terms such as hydrogen bond donor, acceptor, rings, cation, anion, and structural. The TanimotoCombo (Aligned) variant used in this study attempts to minimize the TanimotoCombo between the two ligands by altering the position, but not the conformation, of the molecules, which represents what we use prospectively.

### Bemis-Murcko Scaffold Clustering

The generic scaffold as implemented by RDKit was generated for all ligands from the Moon-shot dataset, and all ligands sharing the same scaffold are clustered together.

#### Scoring

Docking results were scored using OpenEye’s POSIT and Chemgauss4 scores, as well as the RMSD of the docked structure to the known structure.

#### POSIT Score

The OpenEye POSIT score is the estimated probability that the pose of the molecule is *≤* 2 Å RMSD to the known pose, given that the ligand binds the target. It is not intended to predict binding affinity as a comparison between different ligands, but rather identify the best pose for the ligand.

#### Chemgauss4 Score

Chemgauss3 uses Gaussian smoothed potentials of the following interactions: shape, hydrogen bonding between ligand and protein, hydrogen bonding with implicit solvent (i.e. in aqueous phase), and metal-chelator interactions. The Chemgauss4 score is a modification of the Chemgauss3 scoring function with improved hydrogen bonding terms. The Chemgauss4 score is meant to correlate with binding affinity (although, like most docking scores, usually does not).

#### RMSD Calculation

All RMSDs presented herein were calculated using the OpenEye toolkit’s *OERMSD* function on the ligand heavy (non-hydrogen) atoms, which takes into account automorphisms— symmetrical permutations of the ligand configuration which, if not taken into account, resulting in incorrectly high RMSD for chemically equivalent poses.^103^

#### Success Rate and Bootstrapping

The Success Rate, as used in this paper, is calculated by calculating the fraction of query ligands posed *≤* 2 Å to their original crystal pose. When possible, the confidence interval for this value is reported by performing this calculation 1000 times, using randomized references and then taking the values at the 2.5% and 97.5% percentiles as the lower and upper CI values. Where there is not an easy way to randomize the references, we use the beta function to get the posterior probability of observing the number of successes and failures.

## Acknowledgement

The authors thank OpenEye Scientific Software for providing a free academic license to the OpenEye Toolkits to the Chodera Lab. The authors also thank the OpenEye support team, in particular Joseph Ulbrich, for answering many obscure questions and helping troubleshoot some edge-case issues, some of which led to some code and documentation changes in the OpenEye Toolkits. The authors are also grateful to Marcus Wieder and David Mobley for helpful feedback and discussion. This work used resources from the High-Performance Computing Group at Memorial Sloan Kettering Cancer Center. The authors are grateful to the MSKCC DigITs and HPC team, especially Jamie Cheong, Lohit Valleru, and Monica Chakradeo for their assistance with high-performance computing resources.

## Disclaimer

The content is solely the responsibility of the authors and does not necessarily represent the official views of the National Institutes of Health.

## Supporting Information Available

### Code and data availability

The core docking code is available online on Github (https://github.com/choderalab/drugforge), and the code for analyzing the docking results is in (https://github.com/choderalab/harbor). All code to reproduce the docking analysis and the plots in this manuscript can be found in the Harbor repository (https://github.com/choderalab/sars-cov-2-retro-paper-asap). All analyzed data from this repository is already available as part of the COVID Moonshot Initiative.^74^

## References

(1) Blaney, J. A Very Short History of Structure-Based Design: How Did We Get Here and Where Do We Need to Go? Journal of Computer-Aided Molecular Design 2012, 26, 13–14.

(2) Chen, L.; Morrow, J. K.; Tran, H. T.; Phatak, S. S.; Du-Cuny, L.; Zhang, S. From Laptop to Benchtop to Bedside: Structure-based Drug Design on Protein Targets. Current pharmaceutical design 2012, 18, 1217–1239.

(3) Batool, M.; Ahmad, B.; Choi, S. A Structure-Based Drug Discovery Paradigm. International Journal of Molecular Sciences 2019, 20, 2783.

(4) Burley, S. K.; Wu-Wu, A.; Dutta, S.; Ganesan, S.; Zheng, S. X. F. Impact of Structural Biology and the Protein Data Bank on Us Fda New Drug Approvals of Low Molecular Weight Antineoplastic Agents 2019–2023. Oncogene 2024, 43, 2229–2243.

(5) Borshell, N.; Papp, T.; Congreve, M. Valuation Benefits of Structure-Enabled Drug Discovery. Nature Reviews Drug Discovery 2011, 10, 166–166.

(6) Kuhn, P.; Wilson, K.; Patch, M. G.; Stevens, R. C. The Genesis of High-Throughput Structure-Based Drug Discovery Using Protein Crystallography. Current Opinion in Chemical Biology 2002, 6, 704–710.

(7) Namasivayam, V.; Vanangamudi, M.; Kramer, V. G.; Kurup, S.; Zhan, P.; Liu, X.; Kongsted, J.; Byrareddy, S. N. The Journey of HIV-1 Non-Nucleoside Reverse Transcriptase Inhibitors (NNRTIs) from Lab to Clinic. Journal of Medicinal Chemistry 2019, 62, 4851–4883.

(8) Brown, D. G. An Analysis of Successful Hit-to-Clinical Candidate Pairs. Journal of Medicinal Chemistry 2023, 66, 7101–7139.

(9) Rácz, A.; Mihalovits, L. M.; Beckers, M.; Fechner, N.; Stiefl, N.; Sirockin, F.; McCoull, W.; Evertsson, E.; Lemurell, M.; Makara, G.; Keserű, G. M. The Changing Landscape of Medicinal Chemistry Optimization. Nature Reviews Drug Discovery 2025, 1–18.

(10) Bissantz, C.; Kuhn, B.; Stahl, M. A Medicinal Chemist’s Guide to Molecular Interactions. Journal of Medicinal Chemistry 2010, 53, 5061–5084.

(11) Sabe, V. T.; Ntombela, T.; Jhamba, L. A.; Maguire, G. E. M.; Govender, T.; Naicker, T.; Kruger, H. G. Current Trends in Computer Aided Drug Design and a Highlight of Drugs Discovered via Computational Techniques: A Review. European Journal of Medicinal Chemistry 2021, 224, 113705.

(12) Wang, S.; Wacker, D.; Levit, A.; Che, T.; Betz, R. M.; McCorvy, J. D.; Venkatakrishnan, A. J.; Huang, X.-P.; Dror, R. O.; Shoichet, B. K.; Roth, B. L. D4 Dopamine Receptor High-Resolution Structures Enable the Discovery of Selective Agonists. Science 2017, 358, 381–386.

(13) Bender, B. J.; Gahbauer, S.; Luttens, A.; Lyu, J.; Webb, C. M.; Stein, R. M.; Fink, E. A.; Balius, T. E.; Carlsson, J.; Irwin, J. J.; Shoichet, B. K. A Practical Guide to Large-Scale Docking. Nature Protocols 2021, 16, 4799–4832.

(14) Lyu, J.; Wang, S.; Balius, T. E.; Singh, I.; Levit, A.; Moroz, Y. S.; O’Meara, M. J.; Che, T.; Algaa, E.; Tolmachova, K.; Tolmachev, A. A.; Shoichet, B. K.; Roth, B. L.; Irwin, J. J. Ultra-Large Library Docking for Discovering New Chemotypes. Nature 2019, 566, 224–229.

(15) Cournia, Z.; Allen, B.; Sherman, W. Relative Binding Free Energy Calculations in Drug Discovery: Recent Advances and Practical Considerations. Journal of Chemical Information and Modeling 2017, 57, 2911–2937.

(16) Mey, A. S. J. S.; Allen, B. K.; McDonald, H. E. B.; Chodera, J. D.; Hahn, D. F.; Kuhn, M.; Michel, J.; Mobley, D. L.; Naden, L. N.; Prasad, S.; Rizzi, A.; Scheen, J.; Shirts, M. R.; Tresadern, G.; Xu, H. Best Practices for Alchemical Free Energy Calculations [Article v1.0]. Living Journal of Computational Molecular Science 2020, 2, 18378–18378.

(17) Ross, G. A.; Lu, C.; Scarabelli, G.; Albanese, S. K.; Houang, E.; Abel, R.; Harder, E. D.; Wang, L. The Maximal and Current Accuracy of Rigorous Protein-Ligand Binding Free Energy Calculations. Communications Chemistry 2023, 6, 1–12.

(18) Muegge, I.; Hu, Y. Recent Advances in Alchemical Binding Free Energy Calculations for Drug Discovery. ACS Medicinal Chemistry Letters 2023, 14, 244–250.

(19) Baumann, H. M.; Dybeck, E.; McClendon, C. L.; Pickard, F. C. I.; Gapsys, V.; Pérez-Benito, L.; Hahn, D. F.; Tresadern, G.; Mathiowetz, A. M.; Mobley, D. L. Broadening the Scope of Binding Free Energy Calculations Using a Separated Topologies Approach. Journal of Chemical Theory and Computation 2023, 19, 5058–5076.

(20) Chen, L.; Wu, Y.; Wu, C.; Silveira, A.; Sherman, W.; Xu, H.; Gallicchio, E. Performance and Analysis of the Alchemical Transfer Method for Binding-Free-Energy Predictions of Diverse Ligands. Journal of Chemical Information and Modeling 2024, 64, 250–264.

(21) Ivanenkov, Y.; Zagribelnyy, B.; Malyshev, A.; Evteev, S.; Terentiev, V.; Kamya, P.; Bezrukov, D.; Aliper, A.; Ren, F.; Zhavoronkov, A. The Hitchhiker’s Guide to Deep Learning Driven Generative Chemistry. ACS Medicinal Chemistry Letters 2023, 14, 901–915.

(22) Loeffler, H. H.; He, J.; Tibo, A.; Janet, J. P.; Voronov, A.; Mervin, L. H.; Engkvist, O. Reinvent 4: Modern AI–Driven Generative Molecule Design. Journal of Cheminformatics 2024, 16, 20.

(23) Klarich, K.; Goldman, B.; Kramer, T.; Riley, P.; Walters, W. P. Thompson Sampling-An Efficient Method for Searching Ultralarge Synthesis on Demand Databases. Journal of Chemical Information and Modeling 2024, 64, 1158–1171.

(24) Fromer, J. C.; Coley, C. W. An Algorithmic Framework for Synthetic Cost-Aware Decision Making in Molecular Design. 2024; http://arxiv.org/abs/2311.02187.

(25) Jumper, J. et al. Highly Accurate Protein Structure Prediction with AlphaFold. Nature 2021, 596, 583–589.

(26) Ahdritz, G. et al. OpenFold: Retraining AlphaFold2 Yields New Insights into Its Learning Mechanisms and Capacity for Generalization. Nature Methods 2024, 21, 1514–1524.

(27) Baek, M. et al. Accurate Prediction of Protein Structures and Interactions Using a Three-Track Neural Network. Science 2021, 373, 871–876.

(28) Press Release: The Nobel Prize in Chemistry 2024. https://www.nobelprize.org/prizes/chemistry/2024/press-release/.

(29) Zhang, S.; Zubatyuk, R.; Yang, Y.; Roitberg, A.; Isayev, O. ANI-1xBB: An ANI-Based Reactive Potential for Small Organic Molecules. Journal of Chemical Theory and Computation 2025, 21, 4365–4374.

(30) Wang, Y.; Fass, J.; Kaminow, B.; Herr, J. E.; Rufa, D.; Zhang, I.; Pulido, I.; Henry, M.; Macdonald, H. E. B.; Takaba, K.; Chodera, J. D. End-to-End Differentiable Construction of Molecular Mechanics Force Fields. Chemical Science 2022, 13, 12016–12033.

(31) Takaba, K.; Friedman, A. J.; Cavender, C. E.; Behara, P. K.; Pulido, I.; Henry, M. M.; MacDermott-Opeskin, H.; Iacovella, C. R.; Nagle, A. M.; Payne, A. M.; Shirts, M. R.; Mobley, D. L.; Chodera, J. D.; Wang, Y. Machine-Learned Molecular Mechanics Force Fields from Large-Scale Quantum Chemical Data. Chemical Science 2024, 15, 12861–12878.

(32) Khalak, Y.; Tresadern, G.; Hahn, D. F.; de Groot, B. L.; Gapsys, V. Chemical Space Exploration with Active Learning and Alchemical Free Energies. Journal of Chemical Theory and Computation 2022, 18, 6259–6270.

(33) Gusev, F.; Gutkin, E.; Kurnikova, M. G.; Isayev, O. Active Learning Guided Drug Design Lead Optimization Based on Relative Binding Free Energy Modeling. Journal of Chemical Information and Modeling 2023, 63, 583–594.

(34) Graff, D. E.; Aldeghi, M.; Morrone, J. A.; Jordan, K. E.; Pyzer-Knapp, E. O.; Coley, C. W. Self-Focusing Virtual Screening with Active Design Space Pruning. Journal of Chemical Information and Modeling 2022, 62, 3854–3862.

(35) Zhang, Y.; Vass, M.; Shi, D.; Abualrous, E.; Chambers, J. M.; Chopra, N.; Higgs, C.; Kasavajhala, K.; Li, H.; Nandekar, P.; Sato, H.; Miller, E. B.; Repasky, M. P.; Jerome, S. V. Benchmarking Refined and Unrefined AlphaFold2 Structures for Hit Discovery. Journal of Chemical Information and Modeling 2023, 63, 1656–1667.

(36) Lyu, J. et al. AlphaFold2 Structures Template Ligand Discovery; Preprint, 2023.

(37) Rianjongdee, F.; Scheen, J.; Macdonald, H. B.; Gowers, R.; Degorce, S.; Green, A.; Scully, C.; Duffy, T.; Howes, J.; Cordery, C.; Bucher, A.; Aithani, L.; Domański, J. Leveraging Alchemical Free Energy Calculations with Accurate Protein Structure Prediction. 2025; https://chemrxiv.org/engage/chemrxiv/article-details/67b705ad81d2151a02442c4a.

(38) Abramson, J. et al. Accurate Structure Prediction of Biomolecular Interactions with AlphaFold 3. Nature 2024, 630, 493–500.

(39) Passaro, S.; Corso, G.; Wohlwend, J.; Reveiz, M.; Thaler, S.; Somnath, V. R.; Getz, N.; Portnoi, T.; Roy, J.; Stark, H.; Kwabi-Addo, D.; Beaini, D.; Jaakkola, T.; Barzilay, R. Boltz-2: Towards Accurate and Efficient Binding Affinity Prediction. 2025; https://www.biorxiv.org/content/10.1101/2025.06.14.659707v1.

(40) Corley, N. et al. Accelerating Biomolecular Modeling with AtomWorks and RF3. 2025; https://www.biorxiv.org/content/10.1101/2025.08.14.670328v2.

(41) Nittinger, E.; Yoluk, Ö.; Tibo, A.; Olanders, G.; Tyrchan, C. Co-Folding, the Future of Docking – Prediction of Allosteric and Orthosteric Ligands. Artificial Intelligence in the Life Sciences 2025, 8, 100136.

(42) Škrinjar, P.; Eberhardt, J.; Tauriello, G.; Schwede, T.; Durairaj, J. Have Protein-Ligand Cofolding Methods Moved beyond Memorisation? 2025; https://www.biorxiv.org/content/10.1101/2025.02.03.636309v3.

(43) Blundell, T. L.; Jhoti, H.; Abell, C. High-Throughput Crystallography for Lead Discovery in Drug Design. Nature Reviews Drug Discovery 2002, 1, 45–54.

(44) Fearon, D. et al. Accelerating Drug Discovery With High-Throughput Crystallographic Fragment Screening and Structural Enablement. Applied Research 2025, 4, e202400192.

(45) Grosjean, H.; Aimon, A.; Hassell-Hart, S.; Thompson, W.; Koekemoer, L.; Bennett, J.; Anderson, C.; FitzGerald, E. A.; Krojer, T.; Bradley, A.; Fedorov, O.; Biggin, P. C.; Spencer, J.; von Delft, F. High-Throughput Crystallography for Rapid Fragment Growth from Crude Arrays by Low-Cost Robotics. 2023; https://chemrxiv.org/engage/chemrxiv/article-details/641ef81e647e3dca997c2ba6.

(46) Correy, G. J. et al. Extensive Exploration of Structure Activity Relationships for the SARS-CoV-2 Macrodomain from Shape-Based Fragment Merging and Active Learning. 2024; https://www.biorxiv.org/content/10.1101/2024.08.25.609621v1.

(47) Diamond Light Source, Harwell Science and Innovation Campus, Fermi Ave, Didcot OX11 0DE, UK.

(48) XChem @ Diamond. https://fragalysis.diamond.ac.uk/viewer/react/landing.

(49) Saar, K. L. et al. Turning High-Throughput Structural Biology into Predictive Inhibitor Design. Proceedings of the National Academy of Sciences 2023, 120, e2214168120.

(50) Saini, M.; Aschenbrenner, J. C.; Ruiz, F. X.; Chopra, A.; Chandran, A. V.; Marples, P. G.; Balcomb, B. H.; Fearon, D.; von Delft, F.; Arnold, E. Crystallographic Fragment Screening of the Dengue Virus Polymerase Reveals Multiple Binding Sites for the Development of Non-Nucleoside Antiflavivirals. 2025; https://www.biorxiv.org/content/10.1101/2025.03.31.646453v1.

(51) Ni, X. et al. Crystallographic Fragment Screening and Deep Mutational Scanning of Zika Virus NS2B-NS3 Protease Enable Development of Resistance-Resilient Inhibitors. bioRxiv 2025, 2024.04.29.591502.

(52) Godoy, A. S. et al. High-Throughput Crystallographic Fragment Screening of Zika Virus NS3 Helicase. 2024; https://www.biorxiv.org/content/10.1101/2024.04.27.591279v1.

(53) Grosjean, H. et al. Binding-Site Purification of Actives (B-SPA) Enables Efficient Large-Scale Progression of Fragment Hits by Combining Multi-Step Array Synthesis With HT Crystallography. Angewandte Chemie International Edition 2025, 64, e202424373.

(54) Dubianok, Y.; Kumar, A.; Rak, A. In Target Identification and Validation in Drug Discovery: Methods and Protocols; Moll, J., Carotta, S., Eds.; Springer US: New York, NY, 2025; pp 17–49.

(55) Grey, J.; Thompson, D. Challenges and Opportunities for New Protein Crystallization Strategies in Structure-Based Drug Design. Expert opinion on drug discovery 2010, 5, 1039–1045.

(56) Käck, H.; Sjögren, T. Macromolecular Crystallography from an Industrial Perspective – the Impact of Synchrotron Radiation on Structure-Based Drug Discovery. Journal of Synchrotron Radiation 2025, 32, 294–303.

(57) Cappel, D.; Jerome, S.; Hessler, G.; Matter, H. Impact of Different Automated Binding Pose Generation Approaches on Relative Binding Free Energy Simulations. Journal of Chemical Information and Modeling 2020, 60, 1432–1444.

(58) Kuntz, I. D.; Blaney, J. M.; Oatley, S. J.; Langridge, R.; Ferrin, T. E. A Geometric Approach to Macromolecule-Ligand Interactions. Journal of Molecular Biology 1982, 161, 269–288.

(59) Meng, E. C.; Shoichet, B. K.; Kuntz, I. D. Automated Docking with Grid-Based Energy Evaluation. Journal of Computational Chemistry 1992, 13, 505–524.

(60) Jones, G.; Willett, P.; Glen, R. C.; Leach, A. R.; Taylor, R. Development and Validation of a Genetic Algorithm for Flexible Docking1. Journal of Molecular Biology 1997, 267, 727–748.

(61) Friesner, R. A.; Banks, J. L.; Murphy, R. B.; Halgren, T. A.; Klicic, J. J.; Mainz, D. T.; Repasky, M. P.; Knoll, E. H.; Shelley, M.; Perry, J. K.; Shaw, D. E.; Francis, P.; Shenkin, P. S. Glide: A New Approach for Rapid, Accurate Docking and Scoring. 1. Method and Assessment of Docking Accuracy. Journal of Medicinal Chemistry 2004, 47, 1739–1749.

(62) Trott, O.; Olson, A. J. AutoDock Vina: Improving the Speed and Accuracy of Docking with a New Scoring Function, Efficient Optimization, and Multithreading. Journal of Computational Chemistry 2010, 31, 455–461.

(63) Vittorio, S.; Lunghini, F.; Morerio, P.; Gadioli, D.; Orlandini, S.; Silva, P.; Jan Martinovic; Pedretti, A.; Bonanni, D.; Del Bue, A.; Palermo, G.; Vistoli, G.; Beccari, A. R. Addressing Docking Pose Selection with Structure-Based Deep Learning: Recent Advances, Challenges and Opportunities. Computational and Structural Biotechnology Journal 2024, 23, 2141–2151.

(64) McNutt, A. T.; Li, Y.; Meli, R.; Aggarwal, R.; Koes, D. R. GNINA 1.3: The next Increment in Molecular Docking with Deep Learning. Journal of Cheminformatics 2025, 17, 28.

(65) McGann, M. FRED Pose Prediction and Virtual Screening Accuracy. Journal of Chemical Information and Modeling 2011, 51, 578–596.

(66) McGann, M. FRED and HYBRID Docking Performance on Standardized Datasets. Journal of Computer-Aided Molecular Design 2012, 26, 897–906.

(67) Kelley, B. P.; Brown, S. P.; Warren, G. L.; Muchmore, S. W. POSIT: Flexible Shape-Guided Docking For Pose Prediction. Journal of Chemical Information and Modeling 2015, 55, 1771–1780.

(68) Shamsian, S.; Sokouti, B.; Dastmalchi, S. Benchmarking Different Docking Protocols for Predicting the Binding Poses of Ligands Complexed with Cyclooxygenase Enzymes and Screening Chemical Libraries. BioImpacts : BI 2024, 14, 29955.

(69) Hartshorn, M. J.; Verdonk, M. L.; Chessari, G.; Brewerton, S. C.; Mooij, W. T. M.; Mortenson, P. N.; Murray, C. W. Diverse, High-Quality Test Set for the Validation of Protein-Ligand Docking Performance. Journal of Medicinal Chemistry 2007, 50, 726–741.

(70) Tuccinardi, T.; Botta, M.; Giordano, A.; Martinelli, A. Protein Kinases: Docking and Homology Modeling Reliability. Journal of Chemical Information and Modeling 2010, 50, 1432–1441.

(71) Schaller, D. A.; Christ, C. D.; Chodera, J. D.; Volkamer, A. Benchmarking Cross-Docking Strategies in Kinase Drug Discovery. Journal of Chemical Information and Modeling 2024, 64, 8848–8858.

(72) Cleves, A. E.; Jain, A. N. Knowledge-guided docking: accurate prospective prediction of bound configurations of novel ligands using Surflex-Dock. Journal of Computer-Aided Molecular Design 2015, 29, 485–509.

(73) Yang, X.; Liu, Y.; Gan, J.; Xiao, Z.-X.; Cao, Y. FitDock: protein-ligand docking by template fitting. Briefings in Bioinformatics 2022, 23, bbac087.

(74) Boby, M. L.; Fearon, D.; Ferla, M.; Filep, M.; Koekemoer, L.; Robinson, M. C.; THE COVID MOONSHOT CONSORTIUM; Chodera, J. D.; Lee, A. A.; London, N.; von Delft, A.; von Delft, F. Open Science Discovery of Potent Noncovalent SARS-CoV-2 Main Protease Inhibitors. Science 2023, 382, eabo7201.

(75) Douangamath, A. et al. Crystallographic and Electrophilic Fragment Screening of the SARS-CoV-2 Main Protease. Nature Communications 2020, 11, 5047.

(76) von Delft, F.; Calmiano, M.; Chodera, J.; Griffen, E.; Lee, A.; London, N.; Matviuk, T.; Perry, B.; Robinson, M.; von Delft, A. A White-Knuckle Ride of Open COVID Drug Discovery. Nature 2021, 594, 330–332.

(77) Griffen, E. J. et al. Open-Science Discovery of DNDI-6510, a Compound That Addresses Genotoxic and Metabolic Liabilities of the COVID Moonshot SARS-CoV-2 Mpro Lead Inhibitor. 2025; https://www.biorxiv.org/content/10.1101/2025.06.16.660018v1.

(78) Lee, A. A. et al. Discovery of Potent SARS-CoV-2 Nsp3 Macrodomain Inhibitors Uncovers Lack of Translation to Cellular Antiviral Response. bioRxiv 2024, 2024.08.19.608619.

(79) La, V. N. T.; Kang, L.; Minh, D. D. L. Enzyme Kinetics Model for the Coronavirus Main Protease Including Dimerization and Ligand Binding. Biophysical Journal 2025, 124, 2627–2638.

(80) Yang, Y.; Luo, Y.-D.; Zhang, C.-B.; Xiang, Y.; Bai, X.-Y.; Zhang, D.; Fu, Z.-Y.; Hao, R.-B.; Liu, X.-L. Progress in Research on Inhibitors Targeting SARS-CoV-2 Main Protease (Mpro). ACS Omega 2024, 9, 34196–34219.

(81) Lee, J.; Kenward, C.; Worrall, L. J.; Vuckovic, M.; Gentile, F.; Ton, A.-T.; Ng, M.; Cherkasov, A.; Strynadka, N. C. J.; Paetzel, M. X-Ray Crystallographic Characterization of the SARS-CoV-2 Main Protease Polyprotein Cleavage Sites Essential for Viral Processing and Maturation. Nature Communications 2022, 13, 5196.

(82) Zhang, L.; Lin, D.; Sun, X.; Curth, U.; Drosten, C.; Sauerhering, L.; Becker, S.; Rox, K.; Hilgenfeld, R. Crystal Structure of SARS-CoV-2 Main Protease Provides a Basis for Design of Improved *α*-Ketoamide Inhibitors. Science (New York, N.y.) 2020, 368, 409–412.

(83) Cao, Y.; Jiang, T.; Girke, T. A Maximum Common Substructure-Based Algorithm for Searching and Predicting Drug-like Compounds. Bioinformatics 2008, 24, i366–i374.

(84) Rogers, D.; Hahn, M. Extended-Connectivity Fingerprints. Journal of Chemical Information and Modeling 2010, 50, 742–754.

(85) Linker, S. M.; Magarkar, A.; Köfinger, J.; Hummer, G.; Seeliger, D. Fragment Binding Pose Predictions Using Unbiased Simulations and Markov-State Models. Journal of Chemical Theory and Computation 2019, 15, 4974–4981.

(86) Lim, N. M.; Osato, M.; Warren, G. L.; Mobley, D. L. Fragment Pose Prediction Using Non-equilibrium Candidate Monte Carlo and Molecular Dynamics Simulations. Journal of Chemical Theory and Computation 2020, 16, 2778–2794.

(87) Vorreiter, C.; Robaa, D.; Sippl, W. Predicting Fragment Binding Modes Using Customized Lennard-Jones Potentials in Short Molecular Dynamics Simulations. Computational and Structural Biotechnology Journal 2024, 27, 102–116.

(88) Carugo, O. How large B-factors can be in protein crystal structures. BMC Bioinformatics 2018, 19, 61.

(89) Smart, O. S.; Horský, V.; Gore, S.; Svobodová Vařeková, R.; Bendová, V.; Kleywegt, G. J.; Velankar, S. Worldwide Protein Data Bank validation information: usage and trends. Acta Crystallographica Section D: Structural Biology 2018, 74, 237–244.

(90) MacDermott-Opeskin, H. et al. A Computational Community Blind Challenge on PanCoronavirus Drug Discovery Data. 2025; https://chemrxiv.org/engage/chemrxiv/article-details/6878bef4fc5f0acb52a813f5.

(91) Scheen, J.; Rianjongdee, F.; Macdonald, H. B.; Gowers, R.; Degorce, S.; Green, A.; Scully, C.; Duffy, T.; Howes, J.; Cordery, C.; Bucher, A.; Aithani, L.; Domański, J. Leveraging Alchemical Free Energy Calculations with Accurate Protein Structure Prediction. 2025; https://chemrxiv.org/engage/chemrxiv/article-details/67d3ea816dde43c908c21ba4.

(92) Castellanos, M. A.; Payne, A. M.; Scheen, J.; MacDermott-Opeskin, H.; Pulido, I.; Balcomb, B. H.; Griffen, E. J.; Fearon, D.; Barr, H.; Lahav, N.; Cousins, D.; Stacey, J.; Robinson, R.; Lefker, B.; Chodera, J. D. A Structure-Based Computational Pipeline for Broad-Spectrum Antiviral Discovery. 2025; https://www.biorxiv.org/content/10.1101/2025.07.29.667267v1.

(93) Errington, D.; Schneider, C.; Bouysset, C.; Dreyer, F. A. Assessing Interaction Recovery of Predicted Protein-Ligand Poses. Journal of Cheminformatics 2025, 17, 76.

(94) Burley, S. K.; Berman, H. M. Open Access Data: A Cornerstone for Artificial Intelligence Approaches to Protein Structure Prediction. Structure (London, England : 1993) 2021, 29, 515–520.

(95) Unoh, Y. et al. Discovery of S-217622, a Noncovalent Oral SARS-CoV-2 3CL Protease Inhibitor Clinical Candidate for Treating COVID-19. Journal of Medicinal Chemistry 2022, 65, 6499–6512.

(96) Heyndrickx, W. et al. MELLODDY: Cross-pharma Federated Learning at Unprecedented Scale Unlocks Benefits in QSAR without Compromising Proprietary Information. Journal of Chemical Information and Modeling 2024, 64, 2331–2344.

(97) Case Study: Drug Discovery AI Consortium. https://www.apheris.com/resources/learning-hub/article/case_study_aisb.

(98) Reinecke, M.; Brear, P.; Vornholz, L.; Berger, B.-T.; Seefried, F.; Wilhelm, S.; Samaras, P.; Gyenis, L.; Litchfield, D. W.; Médard, G.; Müller, S.; Ruland, J.; Hyvönen, M.; Wilhelm, M.; Kuster, B. Chemical Proteomics Reveals the Target Landscape of 1,000 Kinase Inhibitors. Nature Chemical Biology 2024, 20, 577–585.

(99) Edfeldt, K. et al. A Data Science Roadmap for Open Science Organizations Engaged in Early-Stage Drug Discovery. Nature Communications 2024, 15, 5640.

(100) Edwards, A. M.; Owen, D. R. Protein–Ligand Data at Scale to Support Machine Learning. Nature Reviews Chemistry 2025, 1–12.

(101) OpenBind. https://openbind.uk/.

(102) Young, J. Y. et al. OneDep: Unified wwPDB System for Deposition, Biocuration, and Validation of Macromolecular Structures in the PDB Archive. Structure 2017, 25, 536–545.

(103) Bell, E. W.; Zhang, Y. DockRMSD: An Open-Source Tool for Atom Mapping and RMSD Calculation of Symmetric Molecules through Graph Isomorphism. Journal of Cheminformatics 2019, 11, 40.

