## Supporting Information for "How many crystal structures do you need to trust your docking results?"

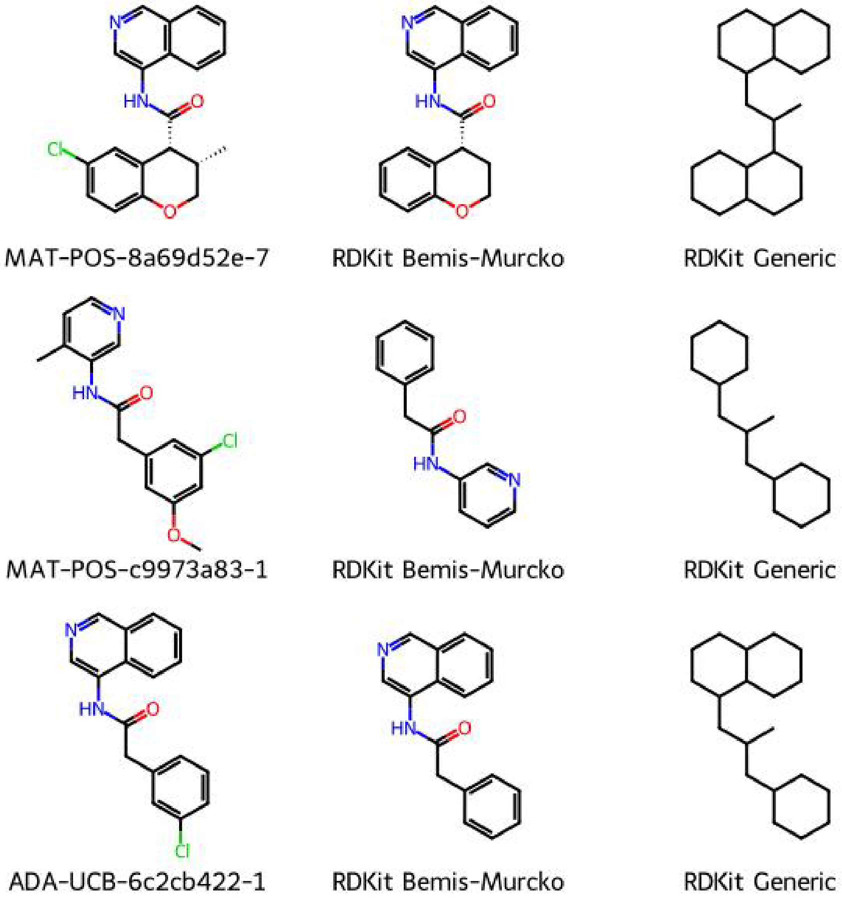


Figure S1: **Many different methods exist to automatically generate scaffolds from ligands.** The Bemis-Murcko scaffold implementations retain more information, but are sensitive to common scaffold modifications like nitrogen-walks. The generic scaffold still retains a chemically intuitive meaning of scaffold while grouping more molecules together.

Table S1: **Pose prediction success rate [95% confidence interval] for each number of same-scaffold reference structures.**

| Scaffold | Ranking | Number of same-scaffold reference structures | | | | | |
|---|---|---|---|---|---|---|---|
| | | 1 | 2 | 5 | 10 | 20 | 50 |
| Scaffold 1 | POSIT Prob. | 0.86 [0.65, 0.97] | 0.93 [0.79, 1.00] | 0.98 [0.92, 1.00] | 0.99 [0.94, 1.00] | 0.99 [0.97, 1.00] | 1.00 [0.98, 1.00] |
| ($N$=62) | RMSD | 0.85 [0.60, 0.97] | 0.96 [0.82, 1.00] | 0.99 [0.97, 1.00] | 1.00 [0.98, 1.00] | 1.00 [1.00, 1.00] | 1.00 [1.00, 1.00] |
| Scaffold 2 | POSIT Prob. | 0.72 [0.19, 0.90] | 0.82 [0.59, 0.91] | 0.88 [0.79, 0.93] | 0.90 [0.84, 0.95] | 0.92 [0.88, 0.97] | 0.98 [0.95, 1.00] |
| ($N$=58) | RMSD | 0.71 [0.19, 0.90] | 0.86 [0.71, 0.93] | 0.93 [0.90, 0.97] | 0.96 [0.91, 1.00] | 0.98 [0.95, 1.00] | 1.00 [1.00, 1.00] |
| Scaffold 3 | POSIT Prob. | 0.82 [0.02, 0.93] | 0.89 [0.71, 0.93] | 0.90 [0.84, 0.95] | 0.91 [0.87, 0.95] | 0.92 [0.87, 0.96] | 0.97 [0.95, 1.00] |
| ($N$=55) | RMSD | 0.82 [0.02, 0.93] | 0.92 [0.87, 0.95] | 0.94 [0.93, 0.96] | 0.95 [0.93, 0.98] | 0.96 [0.95, 1.00] | 0.99 [0.98, 1.00] |
| Scaffold 4 | POSIT Prob. | 0.19 [0.10, 0.40] | 0.35 [0.20, 0.60] | 0.68 [0.50, 0.90] | 1.00 [1.00, 1.00] | — | — |
| ($N$=10) | RMSD | 0.19 [0.10, 0.40] | 0.35 [0.20, 0.60] | 0.71 [0.60, 0.90] | 1.00 [1.00, 1.00] | — | — |

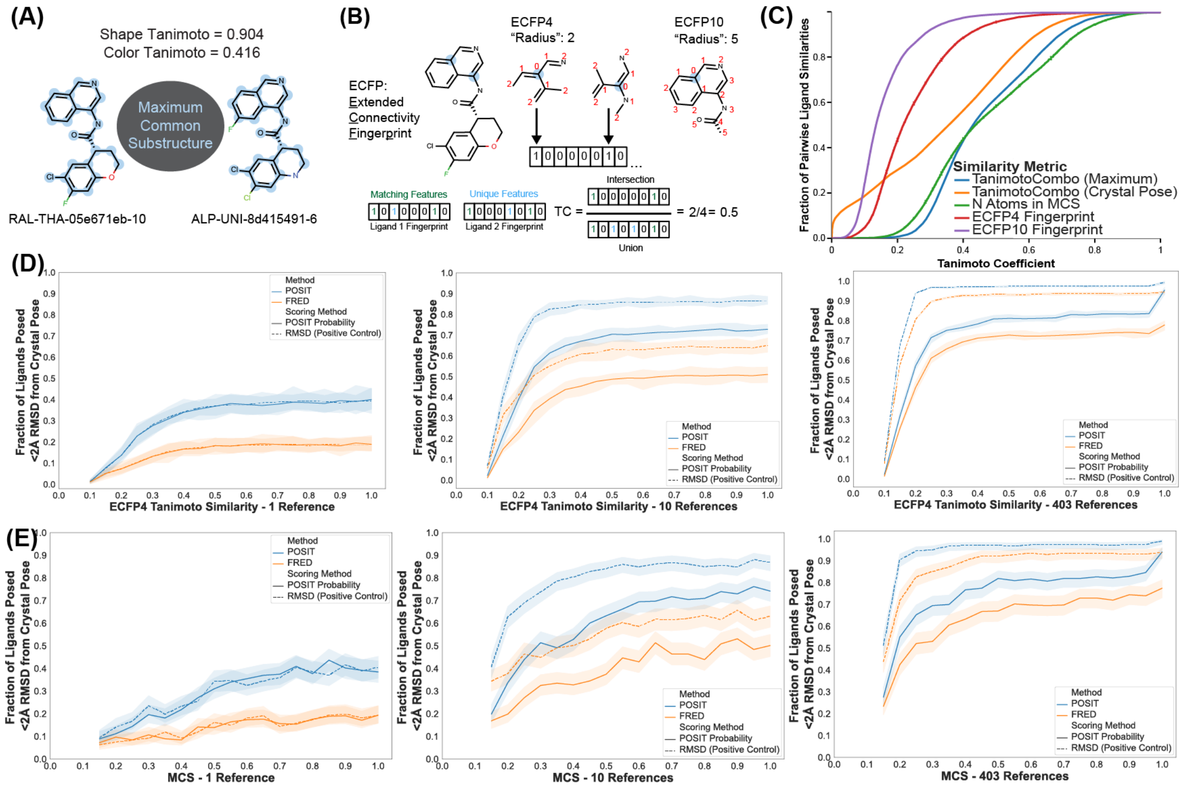


Figure S2: **Different common measurements of ligand similarity have differential impacts on analysis results.** (A) Maximum common substructure (MCS) and (B) Extended Connectivity Fingerprint (ECFP) are other common ways to compare ligand similarity. The Shape and Color Tanimoto scores for two example ligands from the COVID Moonshot are also shown for comparison. (C) The empirical cumulative distribution function plot of all pairwise Tanimoto Coefficients for each of the ligand similarity metrics shows how the choice of similarity metric can change the perception of how chemically diverse a ligand set is. The higher resolution ECFP10 suggests that the COVID Moonshot ligands are quite diverse, with >90% of the pairwise similarities <0.3, whereas the MCS and TanimotoCombo scores suggest that many of the ligands are quite similar. (D) The ECFP4 Tanimoto Similarity does not provide a useful description of this dataset, with most pairwise similarities between 0.1 and 0.4, causing the success rate to quickly plateau. (E) Although the MCS Tanimoto has a similar distribution to the TanimotoCombo, it also plateaus after about 0.5.

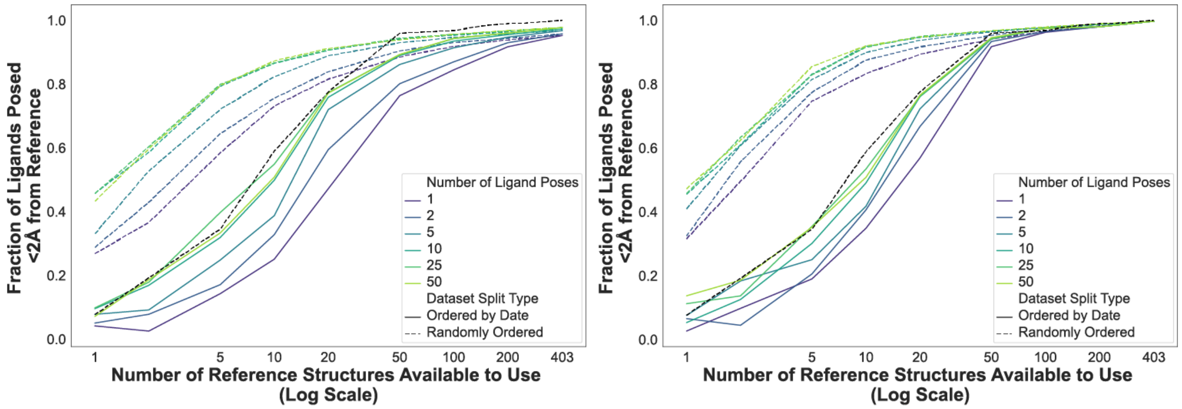


Figure S3: **Improved POSIT settings increase success rate comparable to RMSD ranking of the 50 poses.** The single pose results presented for most of the paper were run with the flags '–allow-retries' and '–relax-mode clash'. For the multipose analysis, these flags were removed in order to save time and compute. The figures are the same from Figure ?? with the addition of a black dotted line representing the Date Split, RMSD-scored POSIT results returning a single pose.

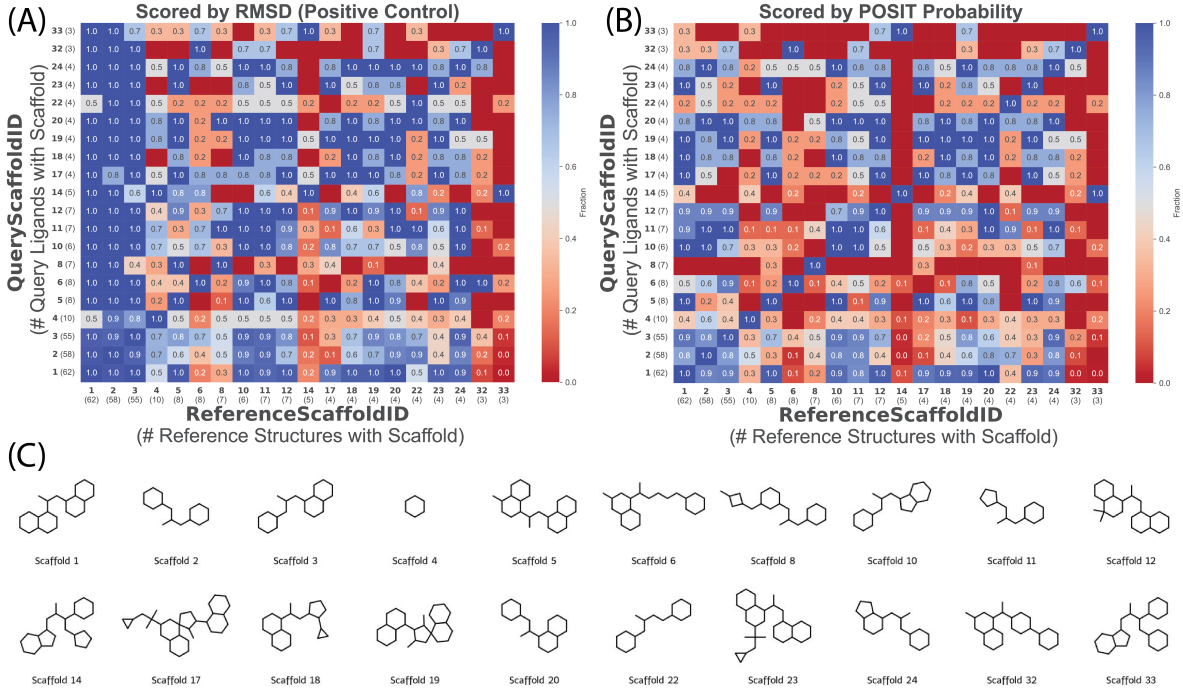


Figure S4: **Scaffold to scaffold cross-docking results analysis.** Heatmap of the fraction of ligands posed within 2Å of their crystal pose for the top 20 most represented scaffolds cross-docked (A) and (B). Each value represents the success rate of the query scaffold (each row) docked to the reference scaffold (each column). On the left (A) the poses are ranked by RMSD, and on the right (B) the poses are ranked by the POSIT Probability. C) The top 20 most represented scaffolds are generic Bemis-Murcko scaffolds are shown below.

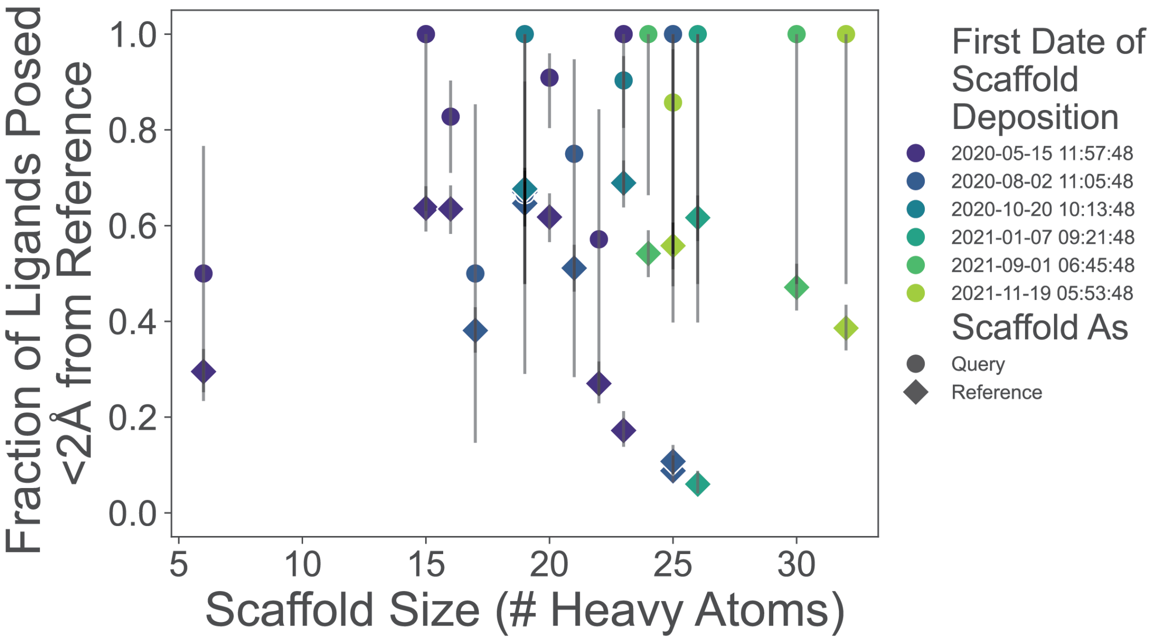


Figure S5: **Pose prediction success does not depend on ligand size above the fragment regime, and ligands were easier to pose than to use as reference.** Results from SI Figure S4 for the top 20 most represented scaffolds averaged for each scaffold, split by either using the scaffolds as query (circle) or reference (diamond). The data points are colored by the date on which the first structure for that scaffold was collected and error bars from bootstrapping over the available references are shown. In all cases, the performance of using a scaffold as the query performed better than using it as a reference.

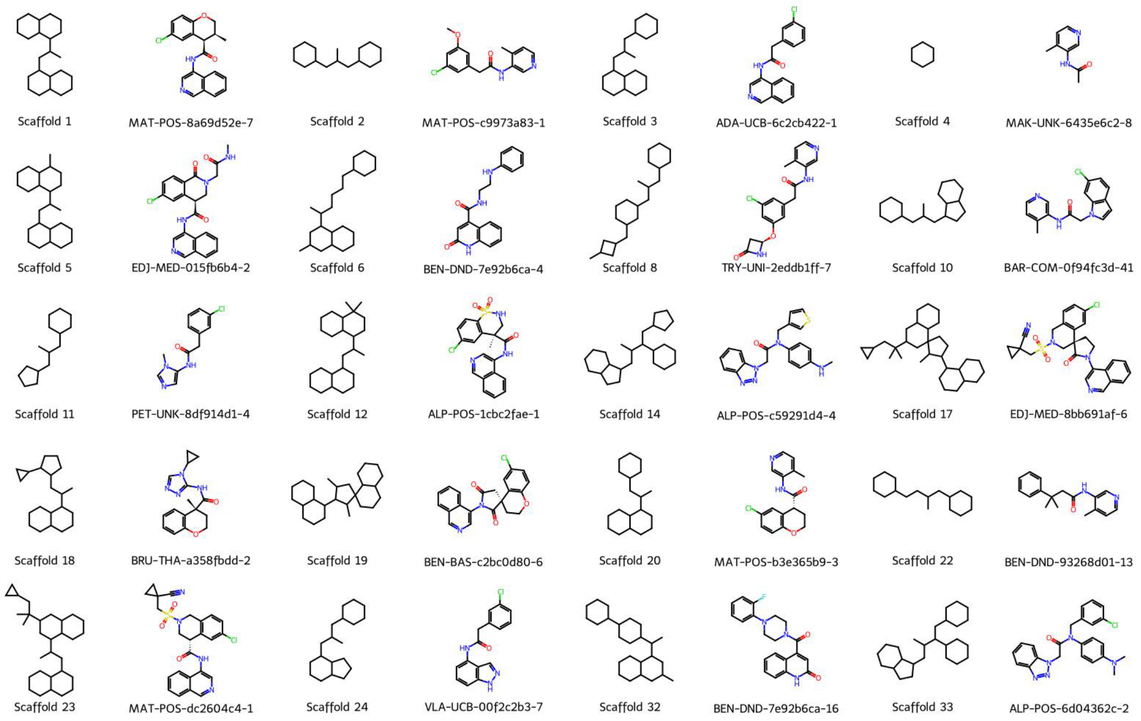


Figure S6: **Top 20 generic Bemis-Murcko scaffolds with examples of ligands from the COVID Moonshot.** The top 20 most represented scaffolds (left) used for SI Figure (S4-S5) with representative examples of these scaffolds (right) pulled from the COVID Moonshot dataset. The generic Bemis-Murcko scaffold captures the core graph of the molecule while remove the heteroatom and bonding information. Several scaffolds are variations of one another—Scaffold 1 is only slightly modified in Scaffolds 5, 12, and 23.

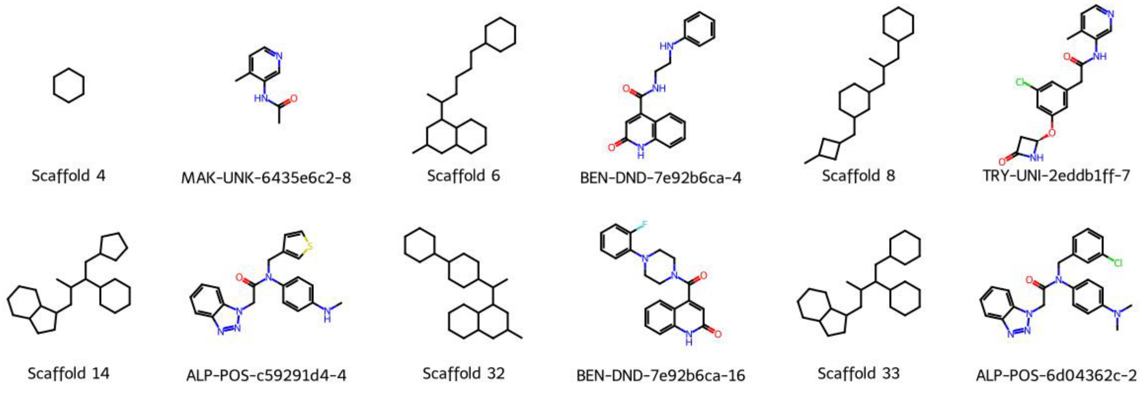


Figure S7: **Challenging scaffolds for pose prediction with examples of ligands from the COVID Moonshot.** The six worst-performing scaffolds (left) from the top 20 are shown alongside representative examples from the COVID Moonshot dataset (right). The fragment scaffold (Scaffold 4) was challenging both to dock and to use as a reference throughout this analysis.

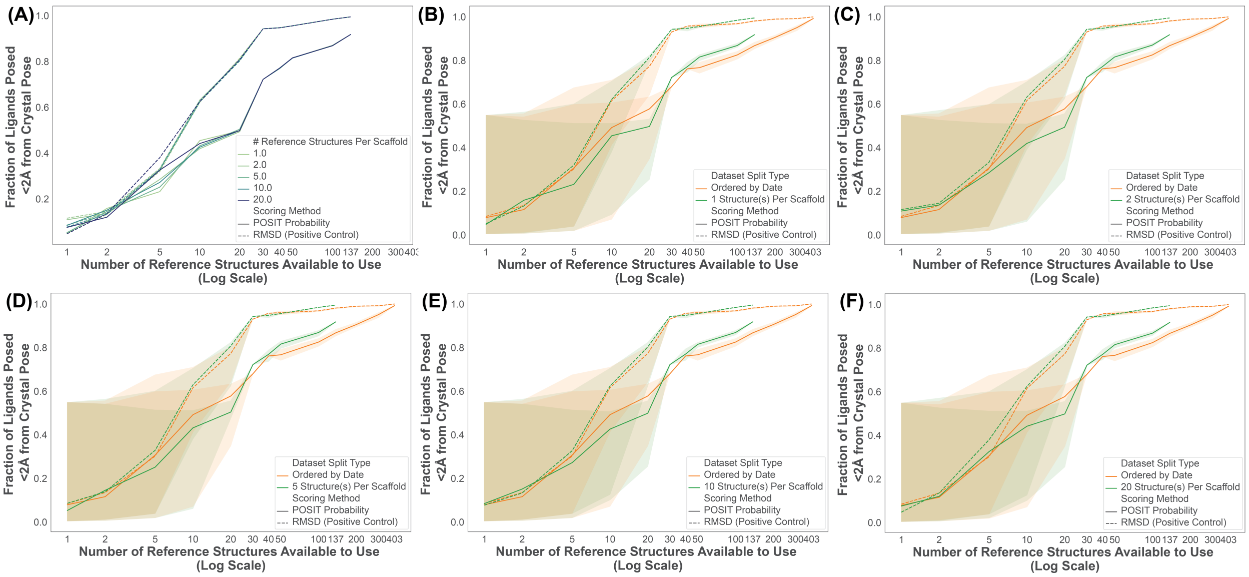


Figure S8: **Increasing the number of structures we collect per scaffold does not show any significant improvement in pose-prediction performance.** (A) The pose prediction performance, reported as the fraction of ligands posed $\leq 2$ Åfrom the crystal pose, is plotted as a function of increasing the number of reference structures available to use, from 1 structure (light green) to 20 (dark blue) for either ranking by the POSIT Probability (solid) or RMSD (dashed). The results 1 structure per scaffold (B) to 20 structures per scaffold (F) in A are compared to the Temporal Split (orange).

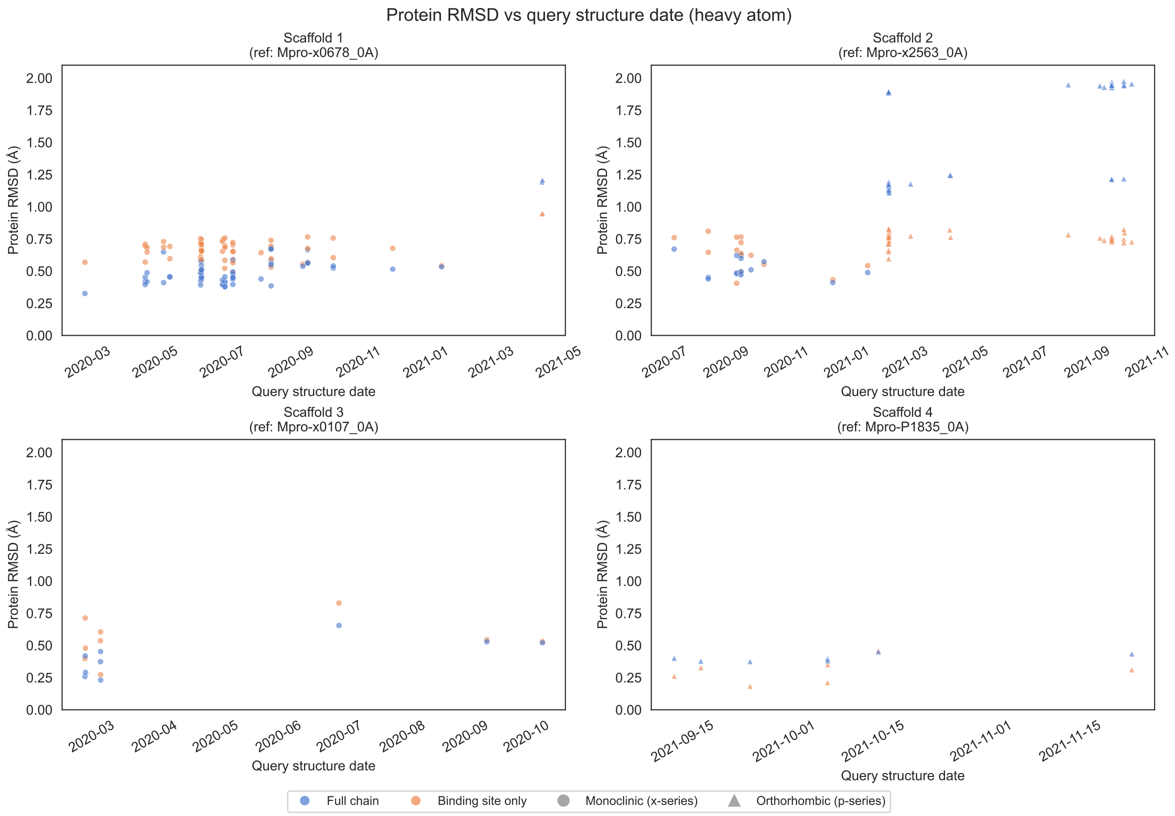


Figure S9: **The SARS-CoV-2 Mpro structures do not show significant conformational changes over time.** The RMSD of the protein heavy atoms between each structure and the first structure collected for that scaffold is plotted as a function of the date on which the structure was collected. The only significant changes were between the monoclinic (x-series, circles) and orthorhobic (p-series, triangles) crystal forms.

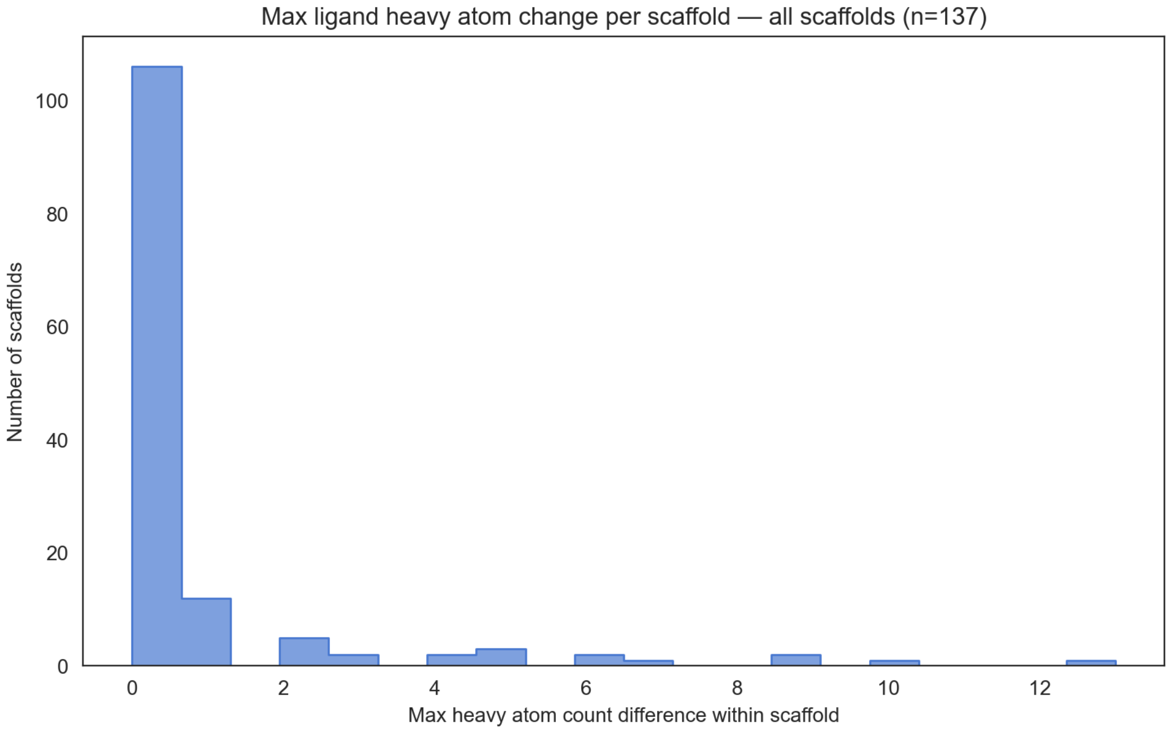


Figure S10: **Within-scaffold chemical excursion across the dataset is modest.** Distribution of the maximum within-scaffold heavy-atom count difference across all 137 generic Bemis-Murcko scaffolds with more than one ligand. The overwhelming majority of scaffolds span at most one heavy atom of variation; the largest excursion observed in the dataset is 13 heavy atoms.

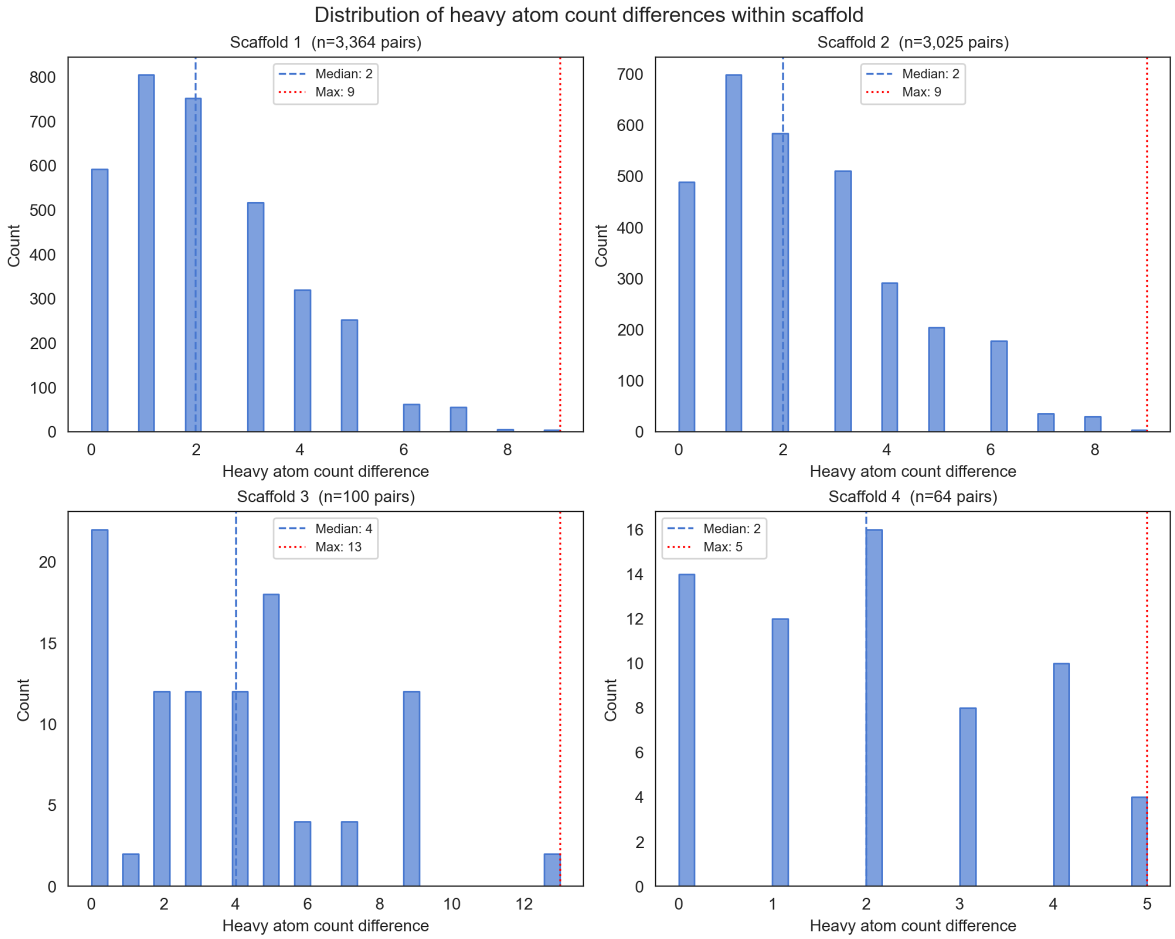


Figure S11: **Pairwise within-scaffold heavy-atom count differences for the top four scaffolds.** Pairwise distributions of heavy-atom count differences within each of the top four scaffolds. Median differences are 2, 2, 4, and 2 heavy atoms for scaffolds 1–4, with maxima of 9, 9, 13, and 5 respectively.

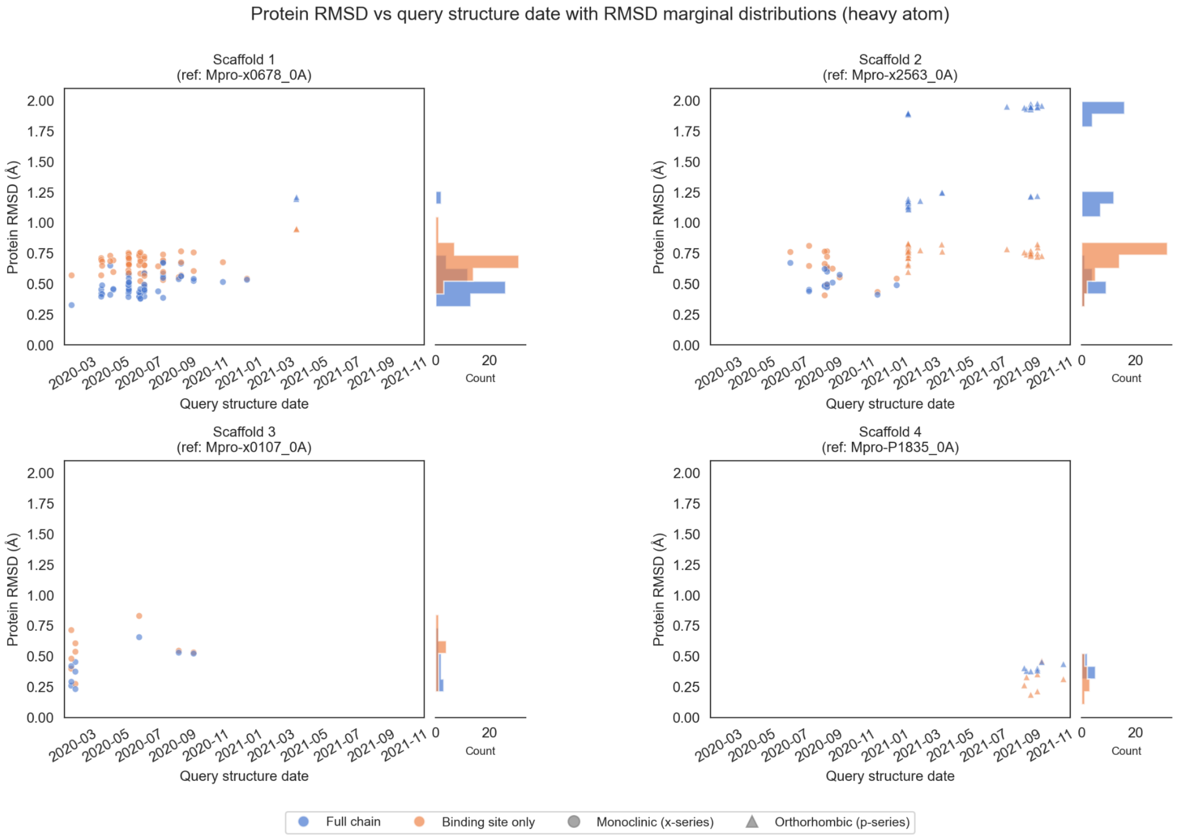


Figure S12: **Protein heavy-atom RMSDs across the four most populous scaffolds are dominated by the crystal-form, not by ligand-induced rearrangement.** Protein heavy-atom RMSD plotted against query structure date for the four most populous scaffolds, with full-chain RMSD (blue) and binding-site-only RMSD (orange) shown separately and marginal histograms on the right. Marker shape distinguishes monoclinic (circle) and orthorhombic (triangle) crystal forms. The full-chain RMSD remains below 2 Å and the binding-site RMSD below 1 Å across all four scaffolds; the upper tail of the distribution is dominated by cross-form (monoclinic vs. orthorhombic) comparisons rather than by ligand-induced rearrangement.

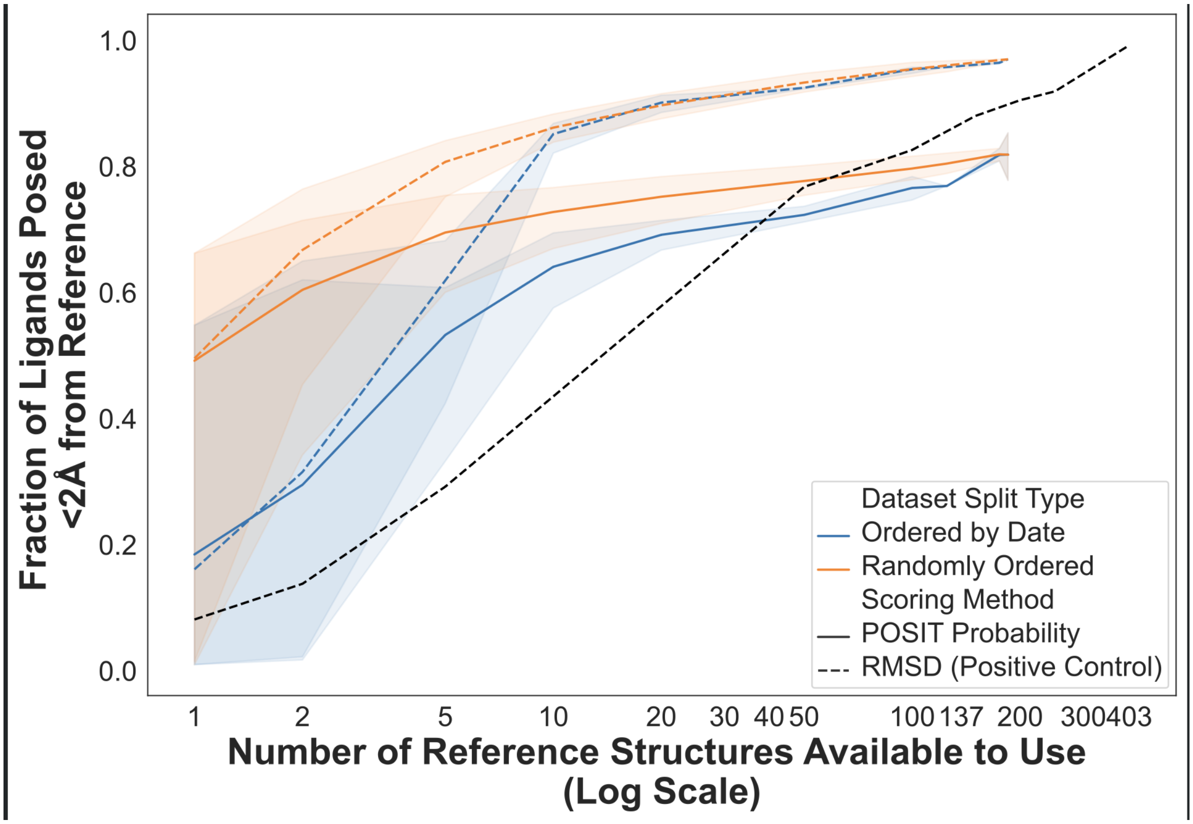


Figure S13: **The Temporal Split underperformance at small reference counts is driven by small early fragments, not by intrinsic time dependence.** Fraction of ligands posed within 2 Å of the crystal pose as a function of the number of reference structures available, for both the Random Split (orange) and the Temporal Split (blue), with both POSIT Probability (solid) and RMSD oracle (dashed) shown alongside. Restricting to ligands with ≥15 heavy atoms brings the early Temporal-Split success rate into close agreement with the Random-Split success rate. This cutoff illustrates the separation from early single-ring fragments of scaffold 4 from the elaborated series; it was not fitted, and no conclusion in this work depends on its precise value.

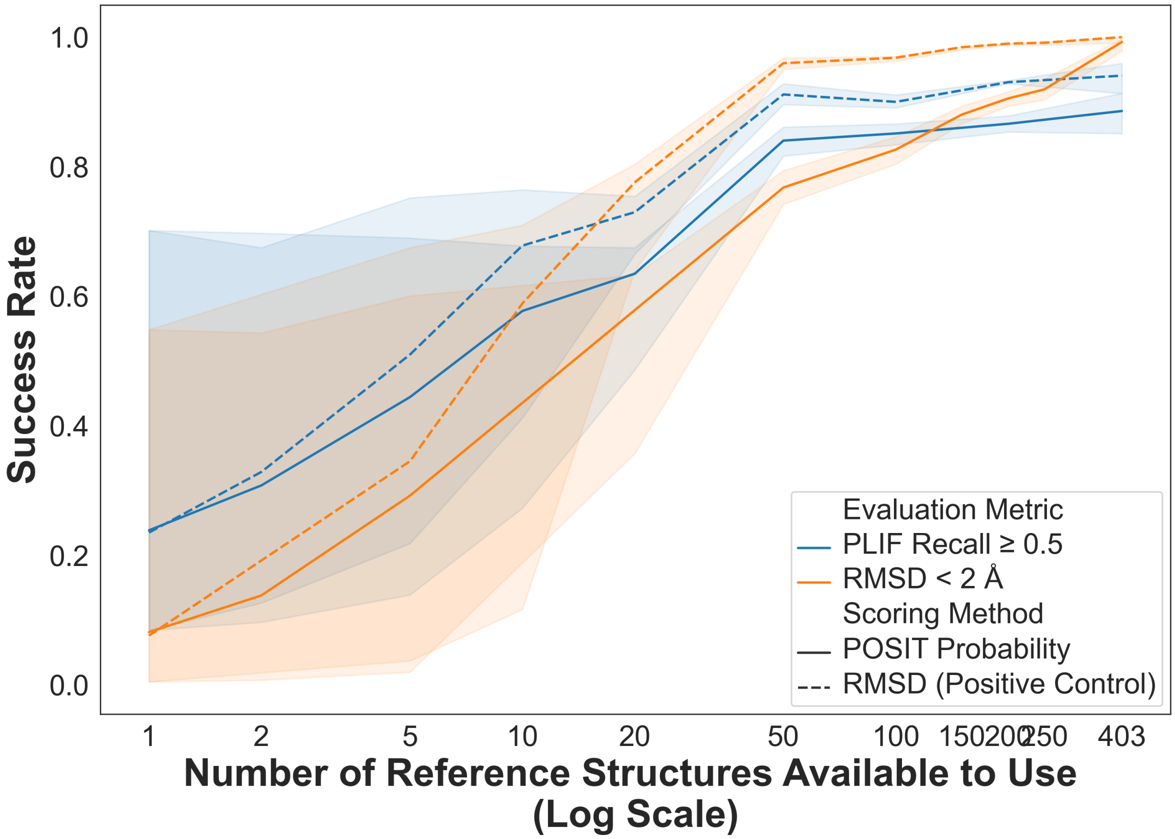


Figure S14: **Re-evaluation of pose prediction success using a PLIF Recall criterion in place of 2 Å heavy-atom RMSD.** Success rate as a function of the number of reference structures available, comparing the PLIF Recall $\geq 0.5$ criterion (blue) against the conventional 2 Å heavy-atom RMSD criterion (orange), with both POSIT Probability (solid) and the RMSD positive control (dashed) shown alongside. The qualitative trends are unchanged: success rates increase with the number of available reference structures and the diminishing-returns plateau is preserved.

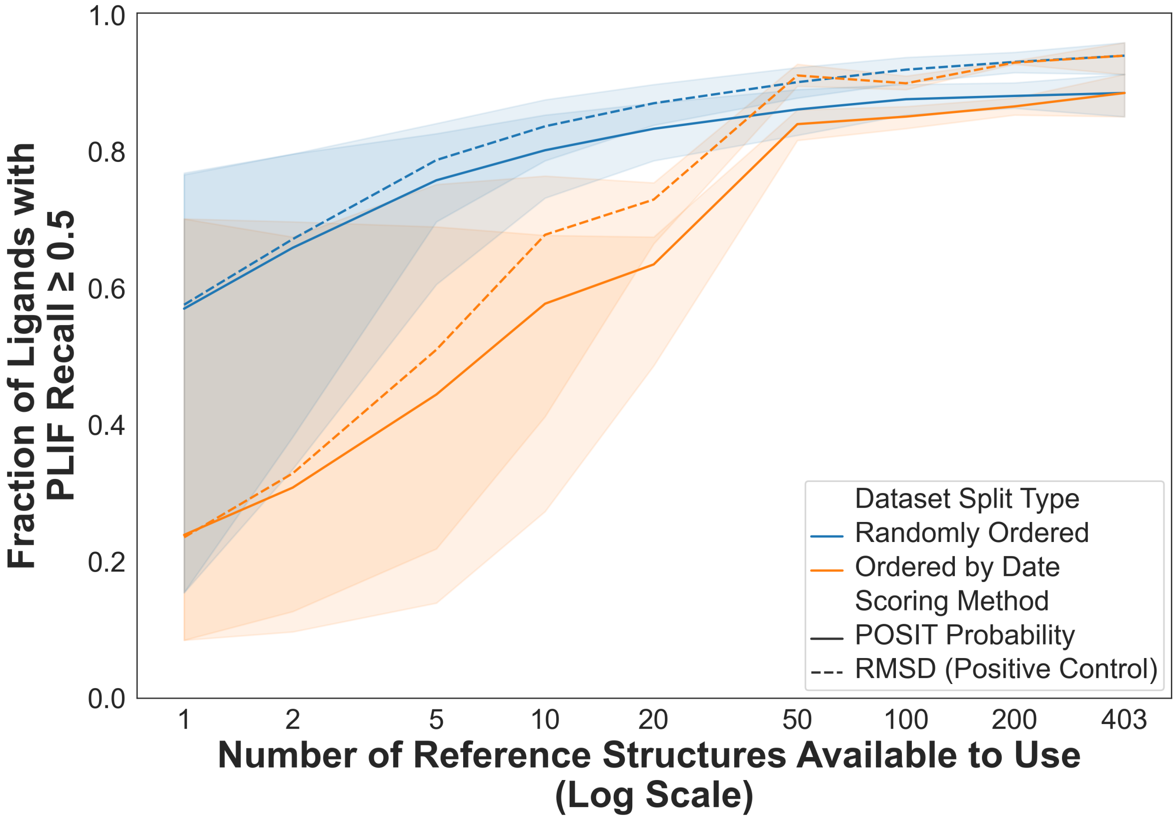


Figure S15: **PLIF Recall success rate as a function of the number of available reference structures, for the Random and Temporal Splits.** Fraction of ligands with PLIF Recall $\geq 0.5$ plotted as a function of the number of reference structures available, for the Random Split (blue) and the Temporal Split (orange), with poses ranked by the POSIT Probability (solid) and by RMSD as a positive control (dashed). The diminishing-returns plateau emerges at the same number of structures as under the 2 Å RMSD criterion (cf. Figure **??**), supporting our central claim that scaffold diversity rather than within-scaffold depth drives most of the gains.

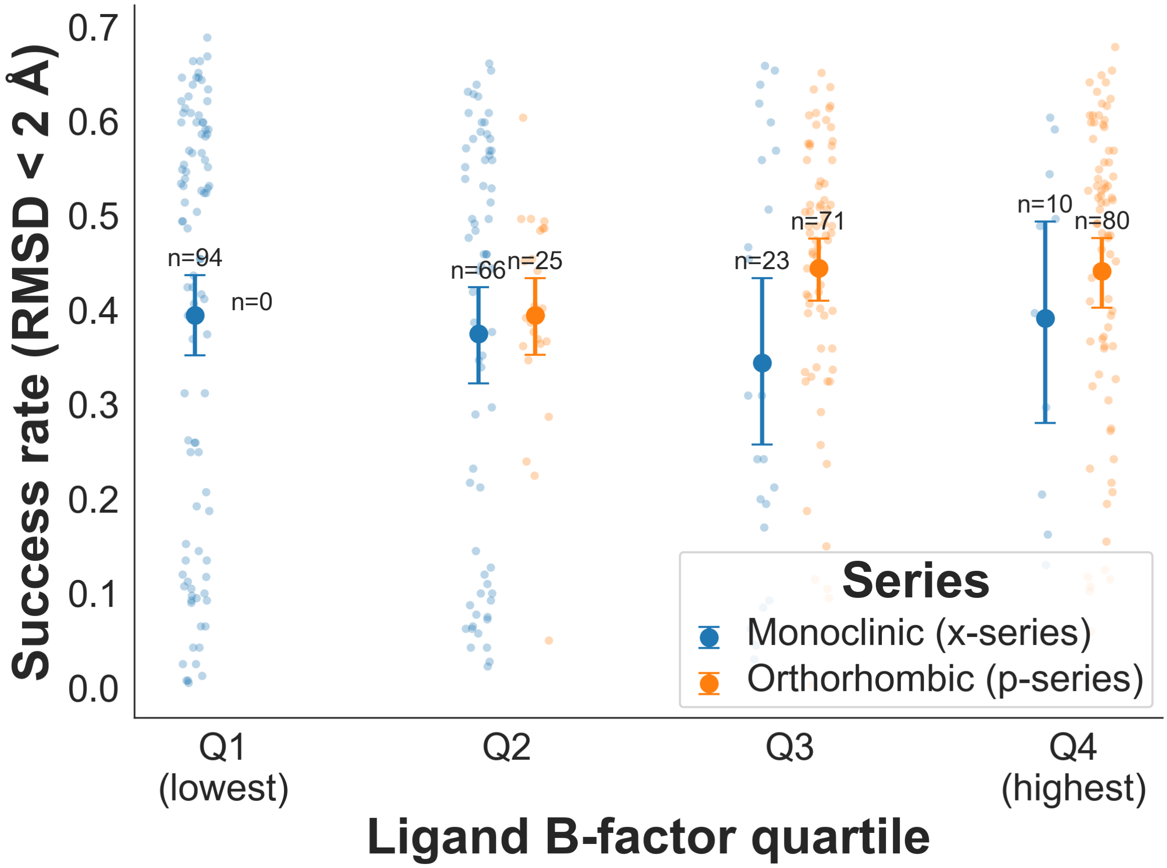


Figure S16: **Per-reference success rate is more strongly affected by crystal form than by ligand B-factor.** Average per-reference success rate (fraction of query ligands posed within 2 Å of the crystal pose, across all query ligands) plotted as a function of the ligand B-factor binned by quartile, separated by crystal form (monoclinic / x-series in blue, orthorhombic / p-series in orange). Despite the higher (worse) ligand B-factors of the orthorhombic structures, those structures generally outperform the monoclinic ones as references, a trend that also held for resolution and R-factor (Figure S17).

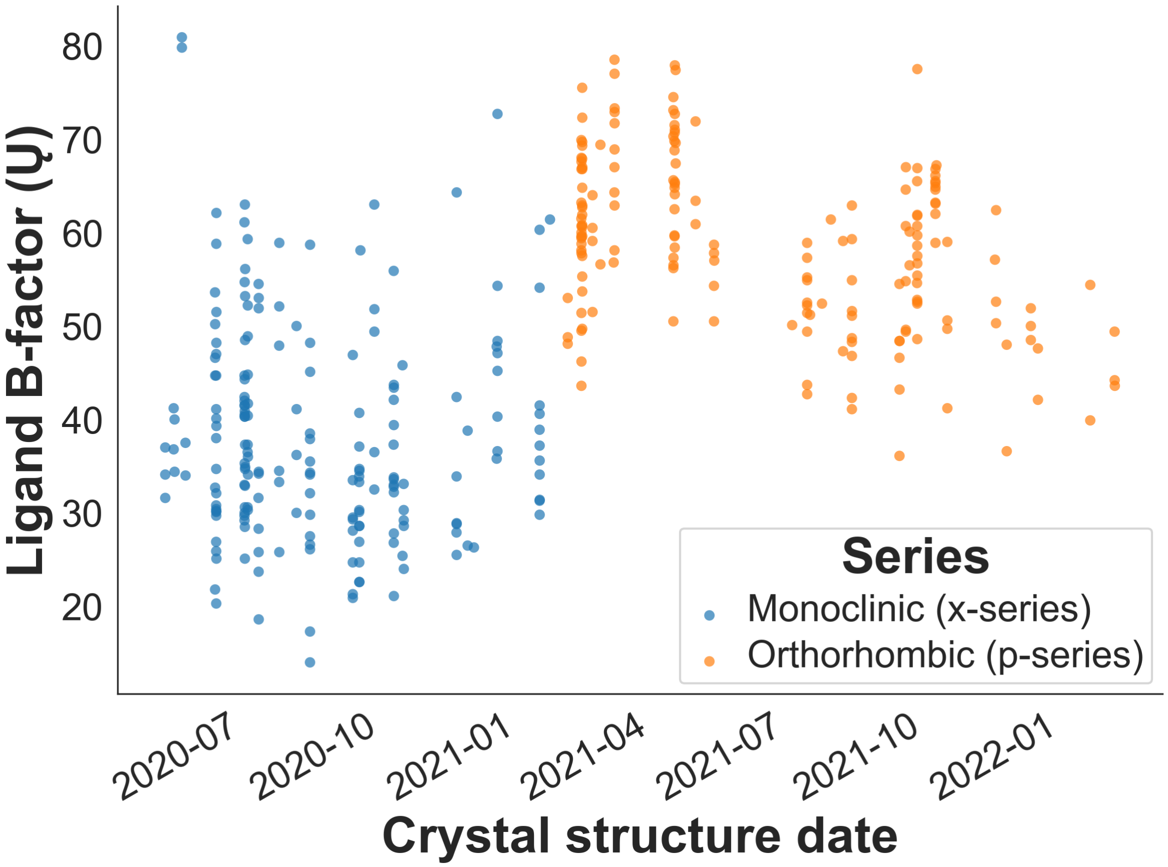


Figure S17: **Ligand B-factors over time, separated by crystal form.** Ligand B-factor plotted against crystal structure collection date, separated by crystal form (monoclinic / x-series in blue, orthorhombic / p-series in orange). The orthorhombic structures, collected later in the campaign, exhibit systematically higher (worse) ligand B-factors than the monoclinic structures, indicating that the better performance of the p-series structures as references (Figure S16) cannot be attributed to better crystal quality alone and is more likely driven by time-dependent chemical similarity trends.

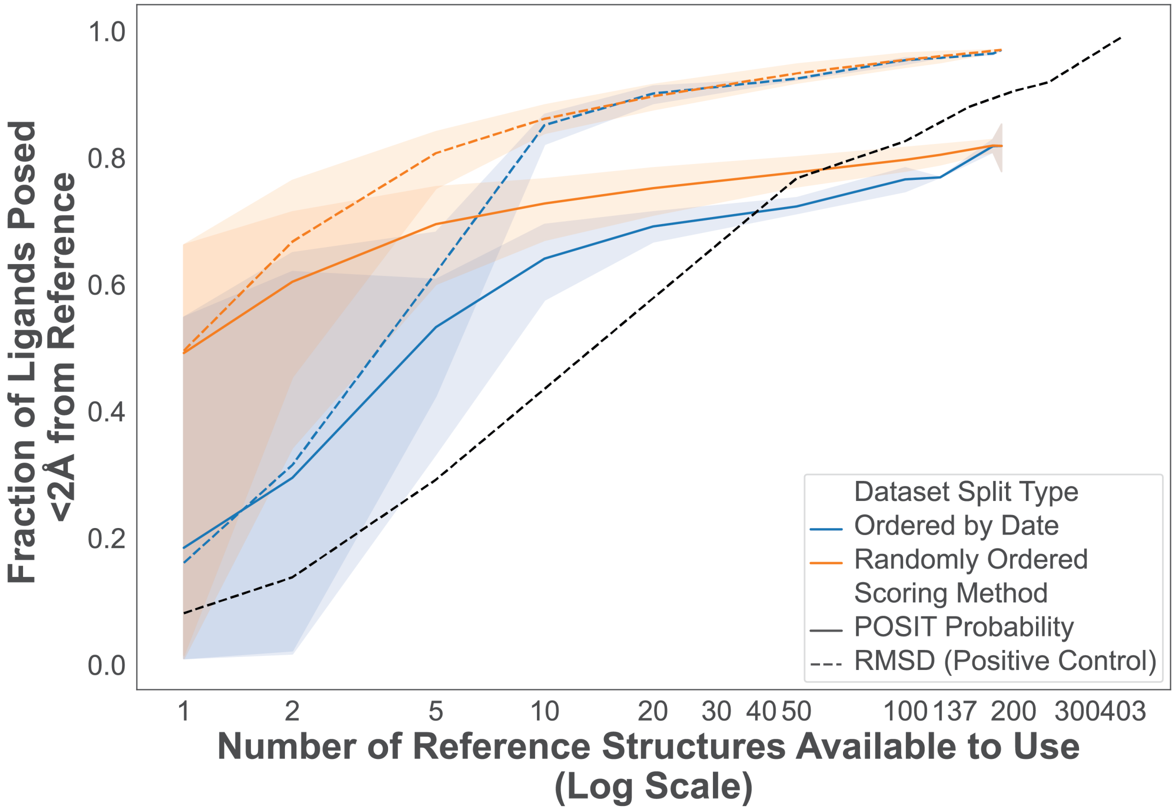


Figure S18: **Pose prediction success for query ligands whose scaffold is not among the top four, docked only to reference structures bearing the top four scaffolds.** This figure supports the statement in the Results that a success rate of 60% is reached at ten structures and had been referenced in the main text without being included in the Supporting Information.

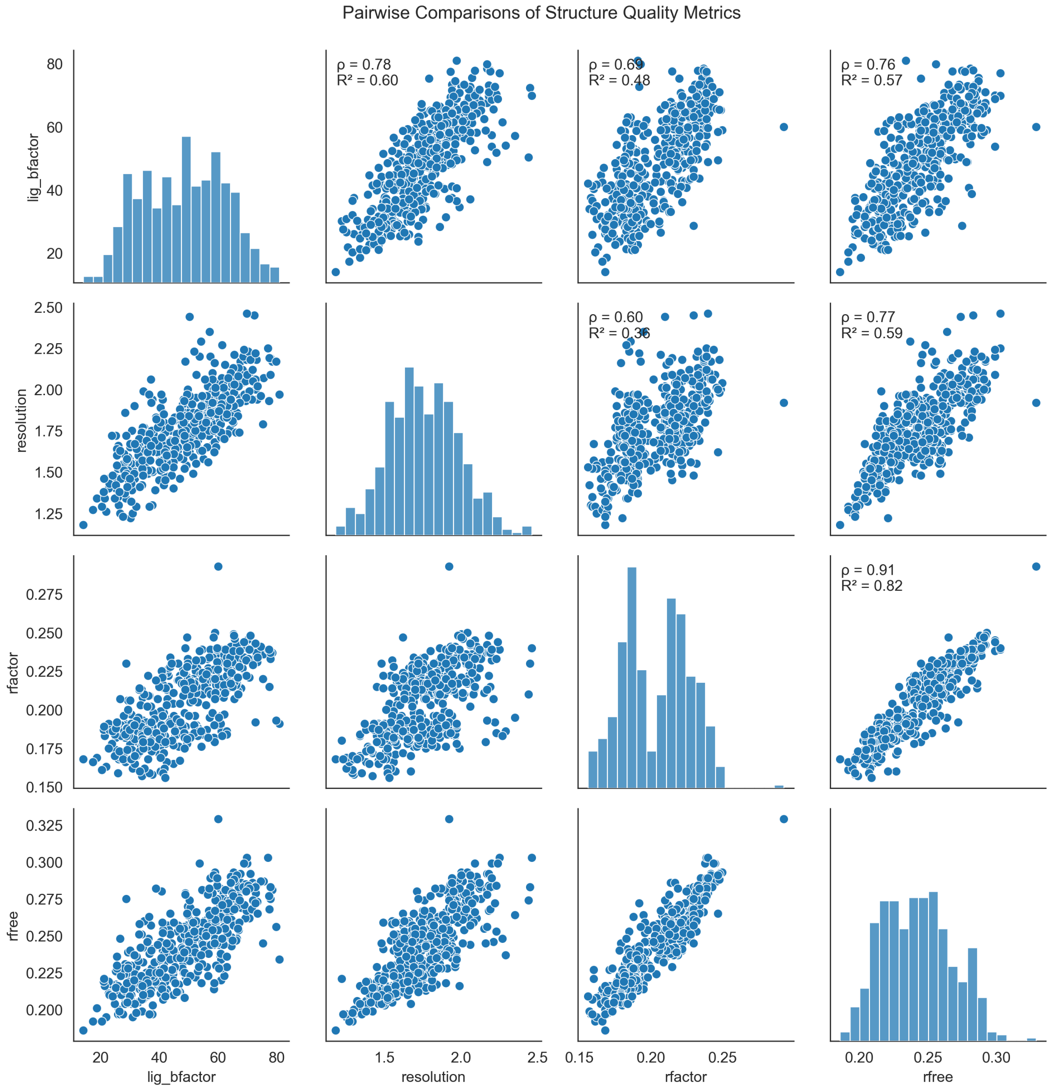


Figure S19: **Standard crystallographic quality metrics are strongly intercorrelated across the dataset.** Pairwise distributions of mean ligand B-factor, resolution, R-factor and R-free across the 403 prepared structures, with Spearman correlation coefficients annotated.

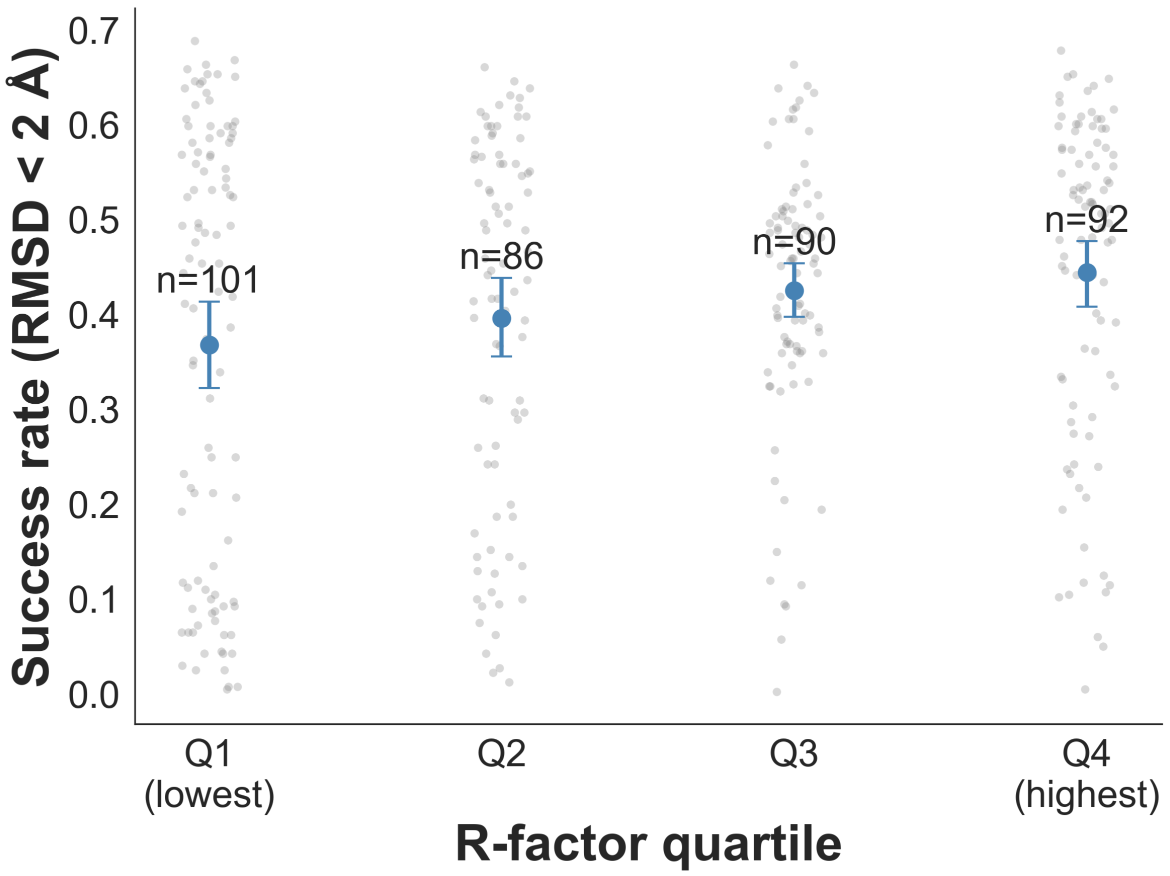


Figure S20: **Per-reference pose prediction success does not degrade as R-factor worsens.** Average per-reference success rate against R-factor quartile, pooled across both crystal forms.
